# Active strains in the basal organ of Corti in gerbil

**DOI:** 10.64898/2026.01.27.702095

**Authors:** Kaitlyn H. Wong, C. Elliott Strimbu, Elizabeth S. Olson

## Abstract

Optical coherence tomography (OCT) has allowed in vivo recording of sound-induced vibrations of different regions within the organ of Corti complex (OCC), including the basilar membrane (BM), outer hair cell/Deiters cell (OHC/DC) region, and reticular lamina (RL). In the hook region of the gerbil cochlea, where measurements can be made with a substantially transverse optical axis, the three regions have different and characteristic motion responses: The OHC/DC region has greater motions than the other two regions at frequencies below the best frequency (sub-BF); the RL region typically has the greatest BF peak and smallest sub-BF motion. The phase of the OHC/DC-region motion increasingly lags BM motion phase as frequency increases; the RL-region motion phase leads BM, but with a relatively small value. All three regions are compressively nonlinear in the BF peak, but only the OHC/DC region shows sub-BF compressive nonlinearity. In this paper, we describe the strain that exists within the RL and OHC-body regions. These strains are large where the motion varies over short distances, and a region of large strain can be as short as a single 2.7 µm measurement pixel, or extend over several pixels, with the extensive strains appearing more often at 70 than at 50 dB SPL. Beyond the region of large strain, over a distance that can exceed 20 μm, the OHC/DC region displays nearly unvarying motion spatially -- this region appears to vibrate as a body.

**Statement of Significance:** The sensory tissue of the cochlea responds actively to a sound stimulus: cell-based forces amplify and enhance the vibration of the sensory tissue. Measurements employing optical coherence tomography have identified major vibration patterns along a sensory-tissue-spanning line that includes the active outer hair cells. In this article, we describe the transitional motion between these major vibration regions and the motion strains that exist as vibration morphs from one region to the next. The findings are presented in frequency response curves to convey the frequency tuning and its stimulus-level dependence, and in one-dimensional heat maps to convey the extent of regional motions and strains. These findings fuel and constrain conceptual and physics-based models of cochlear amplification.

## I. Introduction

In the cochlea, a sound is represented in the pattern of vibration along the long, narrow strip of sensory tissue, the organ of Corti complex (OCC, which refers to cells of the organ of Corti and the acellular basilar membrane (BM) and tectorial membrane (TM) that bound it). To first order, the sound is sorted by frequency with high frequencies represented in the cochlear base and low frequencies in the cochlear apex. In gerbil, the sensory tissue is ∼ 11mm long, with width and thickness about 200 µm x 100 µm, but these vary longitudinally to produce the cochlea’s place-frequency map (1). Fig. 1 shows a labelled schematic and a labelled B-scan of a transverse/radial cross-section of the OCC. The BM is a major structural element of the OCC, composed of radial collagen fibers and a ground substance (2). The TM is a gel-like, collagenous structure containing many specialized proteins (3, 4). The inner and outer pillar cells (PCs) are microtubule-based structural cells that connect the BM to the apex of the OC, the reticular lamina (RL) (5, 6). OHCs are supported by Deiters cells (DCs), which cup the OHCs at their base; this region, containing the nuclei of both cells, is referred to as the OHC/DC region. The DCs contain microtubular structures that extend from the DC feet at the BM to form the DC cup and then extend further as the phalangeal processes, which run from the DC cup to the RL with a transverse/longitudinal slant (7). The RL comprises the apical surfaces of the PCs, the actin-rich termination of DC phalangeal processes, and the cuticular plates of the OHCs and inner hair cells (IHCs) (5). The cuticular plates are composed of a dense matrix of F-actin, one to a few micrometers in thickness. HC stereocilia are anchored into the cuticular plate by rootlets that penetrate deeply into the plate (8–10). In this paper, we describe the dynamic strains that exist in the RL region and the OHC body region due to the motion response to sound stimulation.

**Fig. 1:**
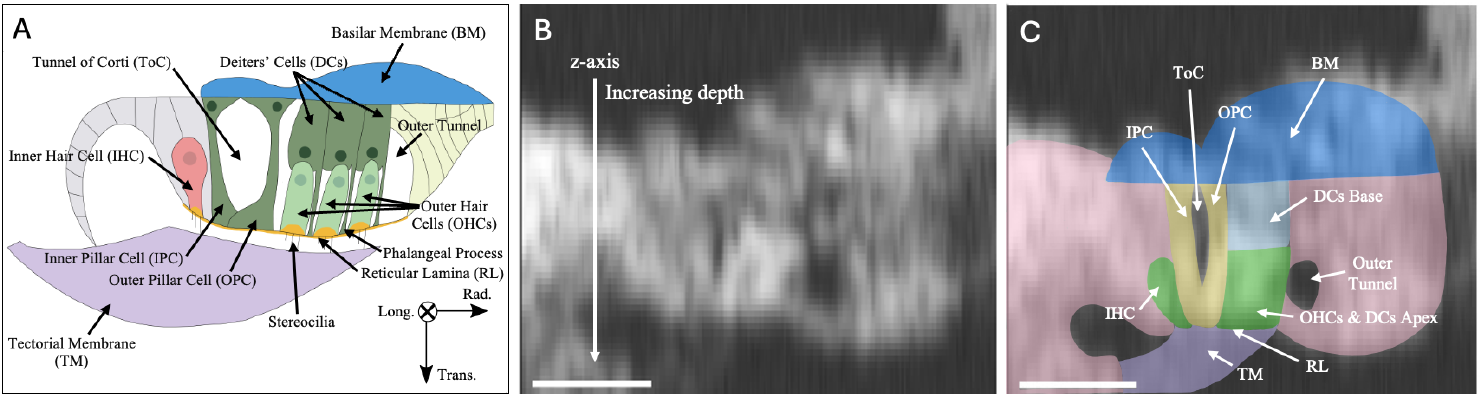
(A) Cartoon diagram of a transverse-radial cross section of the OCC. The diagram includes anatomical axes, longitudinal, radial, and transverse *(l, r, t)*. (B) OCT B-scan of G1047. (C) B-scan from (B) with the relevant regions from (A) approximately delineated. Scale bar = 50 μm. The vertical axis is the optical z-axis, and is not generally aligned with the transverse anatomical axis, although it is shown here that way for simplicity. Because of this B-scan orientation, the radial-transverse cross section shown in OCT-based experiments is typically drawn “upside-down” (as it is here), from its historic orientation, and thus describing the basal and apical ends of the cells can seem upside-down. The base of the PCs and DCs is at the BM, the base of the OHC is held in the DC cup, the apical end of the PCs, HCs, and DCs’ phalangeal processes is at the RL. In reality, the phalangeal processes slant transverse/longitudinal (which could not be indicated in this transverse/radial view, thus here they are shown running purely transverse); Fig. 16 shows the slant.

In the classic description of the cochlear response to sound stimulation, transverse motion of the OCC creates shearing radial motion between the RL and TM, causing HC stereocilia to pivot back and forth, modulating the open probability of the mechanoelectrical transduction (MET) channels at the stereocilia tips, and thus the flow of cations through the HCs. This modulates the transmembrane voltage, and in OHCs activates a piezoelectric protein, prestin, that causes the OHCs to contract and expand, and/or exert contracting/expanding forces (11–14). This OHC action, termed electromotility, is thought to be the fundamental process enhancing the motion of the OCC in a place and frequency dependent manner, which is known as cochlear amplification. Prestin is distributed throughout the OHC basolateral membrane, but not at the cuticular plate (10, 15). The basal pole contains prestin at a reduced density compared to the lateral wall, and charge movement associated with electromotility was measured as greatest at the center of the OHC’s longitudinal extent, with a gradual tapering off towards the apical and basal ends (16). In measurements where the OHC was in a microchamber with one end depolarized and the other hyperpolarized by a voltage applied between the chamber and the bath, one side contracted and the other lengthened (17). Stimulating an OHC with a patch electrode produced a longitudinal strain of up to 1% for 20 mV of quasi-DC stimulation (18), while AC stimulation with a microchamber produced motions of ∼1 nm/mV for OHCs of length ∼50 µm, corresponding to strains of ∼2*10^-5^ / mV (17, 19). A primary goal of the present paper was to observe the quantitative and qualitative character of in vivo sound-induced strains in the OCC, particularly the OHC region.

In past decades, in vivo measurements of sound-evoked motion in the cochlear base were taken solely at the BM, because optical techniques could not penetrate within the sensory tissue. BM motion was tuned at the local best frequency (BF), and compressively nonlinear with input sound pressure level (SPL). Nonlinearity was limited to frequencies near the BF, serving to greatly enhance the BF peak at low-moderate SPL. This observation of tuned amplification was interesting with respect to OHC electromotility, because electromotility as measured in isolated OHCs was broadband (19). With the advent of optical coherence tomography (OCT) (and a related method developed by T. Ren (20)), sound-evoked motions of intra-OCC structures became measurable. One of the first important new findings was that in the OHC region, amplification was broadband, extending sub-BF (20, 21). These boosted, nonlinear sub-BF responses were in keeping with the broadband action of electromotility and could be seen as in vivo evidence of electromotility.

OCT detects motion along the direction of its optical axis. To form a two-dimensional (2D) Brightness-scan (B-scan) image, the optical axis is swept perpendicular to the Axial-scan’s (A-scan) optical-axis, and to form a familiar transverse-radial B-scan image (Fig. 1B), the optical axis should be primarily transverse and the scanning radial. Measurements in the hook region of the gerbil cochlea (∼45 kHz BF) made with a primarily transverse view produced a familiar B-scan image, in which the RL region and faint tectorial membrane (TM) could be identified (22–24). These measurements revealed three basic forms of regional motion: (1) BM displacement peaked and was compressively nonlinear at frequencies around the local BF. (2) The OHC/DC region peaked at the local BF at low/moderate SPL, was low-pass at moderate/high SPL, and showed wideband compressive nonlinearity. An increasing phase lag with frequency re: BM motion accumulated ∼3/4 cycle at frequencies close to the BF. (3) RL motion peaked and was compressively nonlinear at frequencies around the local BF. The peak displacement was larger at the RL than at the BM or OHC/DC region. Sub-BF, RL motion could be smaller than the motion at the BM and display expansive nonlinearity at the highest SPL measured (80 dB). At the RL, the phase typically led the BM by less than 1/4 cycle throughout the frequency range of measurement.

In this contribution, we explore the transitions between the BM, OHC/DC, and RL regions by displaying tuning curves and 1D strain maps from many points within an A-scan that spans the regions. The optical axis was primarily transverse but with a substantial negative (basal-pointing) longitudinal component. To give a brief preview of findings: close to and within the RL region and extending into the OHC body, strains could be large due to the transition between RL-like and OHC-like motion that occurred over just one or two imaging pixels (2.7 μm/pixel). Close to the RL, at frequencies around the BF peak, the strain was in phase with RL displacement. About 15 μm from the RL surface of the OHC region, over an A-scan expanse that could exceed 30 μm, a region of nearly uniform, sub-BF amplified motion was observed. Considering previous studies (25, 26), this was likely the OHC/DC region, swinging approximately perpendicular to the longitudinally slanting phalangeal processes.

## II. Methods

### Animal Preparation

Adult Mongolian gerbils (Meriones unguiculatus) of both sexes were used. The experiments were approved by the Institutional Animal Care and Use Committee (IACUC) of Columbia University. The gerbils were anesthetized with intraperitoneal (IP) injections of ketamine (40 mg/kg) and sodium pentobarbital (40 mg/kg). The gerbils were given the analgesic buprenorphine (0.2 mg/kg) at the start of the surgery and then between six and eight hours after the surgery. Lidocaine (2%) was administered subcutaneously at the start of the surgery as a local anesthetic. To maintain an even plane of anesthesia, supplemental doses of pentobarbital were administered throughout the experiment. A tracheotomy was performed to facilitate breathing, and oxygen was sometimes flowed over the tracheotomy. Temperature was maintained with a temperature-controlled warming blanket supplemented with warming lamps and disposable hand warmers. The head was attached to a two-dimensional (2D) goniometer using dental cement to allow for small adjustments to the head position. The pinna, most of the cartilaginous ear canal (EC), and tissue covering the tympanic bulla were removed. A drop of 38% phosphoric acid gel (Pulpdent, Waterford MA) was placed on the bulla for ∼5 minutes to weaken the bone, and the bulla was gently opened using forceps to view the round window (RW). The cochlear condition was assessed and monitored using 2*f*_1_-*f*_2_ distortion product otoacoustic emissions (DPOAEs) in response to tones *f*_*1*_ and *f*_*2*_ with a ratio of *f*_*2*_ /*f*_*1*_ = 1.2. The tones were presented at 50 and 70 dB SPL. The DPOAE responses were robust and indicative of healthy cochleae (Fig. S1 in Supplemental Information). Once the experiment concluded, the gerbils were euthanized with pentobarbital.

### Acoustic stimulation and recording

Acoustic stimuli were delivered closed-field to the EC using a Radio Shack dynamic speaker. A Sokolich ultrasonic microphone was placed ∼1 mm away from the tympanic membrane and coupled to the speaker tube to measure DPOAEs and EC pressures. Multitone stimuli were generated by a Tucker Davis Technologies System 3, which operated at a 130 kHz sampling rate (130,208.33 Hz). The multitone stimuli used were zwuis tone complexes, comprising 35 approximately equally spaced frequencies chosen so that no harmonics or distortion products overlap with a primary frequency up to the third order (27, 28). The zwuis stimuli were delivered for 1 second. The measurement was made at SPLs ranging from 40 to 80 dB SPL in 10 dB steps.

### OCT

A Thorlabs Telesto 320 spectral domain OCT system with a center wavelength of 1300 nm and a 5X LSM03 objective lens was used to perform imaging and vibrometry. According to the manufacturer, the system has axial and lateral resolutions in water of ∼ 4 and 10 μm, respectively. The axial resolution is set by the light source bandwidth and the lateral resolution by the objective lens. The imaging acquisition process starts with a 1D reflectivity map (the A-scan noted above) along the depth of the preparation, the optical *z*-axis. Adjacent pixels in an A-scan are ∼2.7 μm apart. The OCT system uses scanning mirrors to sweep the optical beam along the *x*- or *y*-axis, producing a 2D reflectivity image, the B-scan. If the OCT system sweeps the optical beam in the third orthogonal direction, it yields a 3D volumetric image.

At the beginning of each experiment, the ThorImage program was used to orient the preparation and identify the regions of interest within the OCC. All measurements were made through the RW to obtain a view of the OCC that was primarily transverse in the hook region, where the BF is ∼45 kHz. A series of A-scans recorded at 130 kHz at the same location with concurrent acoustic stimulation produces a motion scan (M-scan), which simultaneously captures sub-pixel displacements of structures along the A-scan. An objective of the current study was to find spatial displacement variations, and a finely spaced data set was desired. Along an A-scan, all of the pixels within the OCC were selected for vibration analysis. The vibrometry data of each pixel were deemed significant if the amplitude at the stimulus frequency was above the noise floor by at least 2.5 times the standard deviation of the noise. The noise floor was determined from a set of adjacent points in the time-frequency Fourier transform. In G986 from an earlier period of experiments, a subset of A-scan pixels was chosen to analyze for displacement. This resulted in a relatively sparse, but still useful, data set for this study. B-scans were acquired before and after each vibrometry measurement and evaluated to ensure that the position of the preparation had not shifted significantly during the recording. Custom C++ software developed using the Thorlabs Spectral Radar software development kit was used for data collection and initial processing of vibrometry data. Further analysis and post-processing were performed in MATLAB. A 3D volumetric image and the planar approximation algorithm (29) was used to determine the anatomical longitudinal, radial, and transverse *(l,r,t)* components of the optical axis for each experiment; the results are included in the figure captions. (The *(l,r,t)* components form a unit vector; they add in quadrature to 1 (or close to 1 due to rounding). For example, *l,r,t* values of -0.6, 0.1, 0.8 would indicate that the measured motion was the sum of -0.6 times the longitudinal component of motion, 0.1 times the radial component of motion, and 0.8 times the transverse component of motion.)

### Calculations

Eq. 1 was used to calculate strain (ε), where *x*_*deep*_ and *x*_*shallow*_ are the displacements measured at the deeper and shallower pixels, respectively. (Fig. 1B clarifies the depth axis.) *x*_*deep*_ − *x*_*shallow*_ is a complex difference and Δ*x* is the distance between the pixels. Results taken with 70 and 50 dB SPL are shown. 70 dB SPL results are shown because that is the lowest SPL for which data were often available over the full frequency range; results at 50 dB SPL are shown mainly in the BF peak.

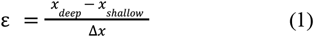

We present the frequency responses in three different color-coded regions (BM, OHC/DC and RL). To defend the separation between groups, the degree of similarity between adjacent response curves was quantified using Eq. 2.

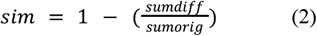

Sumdiff and sumorig are found as follows: sumdiff was found as the magnitude of the frequency-by-frequency complex difference between the two curves, which were then summed across frequency. Sumorig was found as the magnitude values of each of the two original curves summed across frequency, and these two values averaged. If sim was greater than 0.7, by this metric, the two curves were greater than 70% similar and were considered to be within the same region (OHC/DC, RL, or BM), and color-coded green, orange, or blue, respectively. If sim was less than 0.7, the two adjacent curves were considered to be in different groups. The fraction (sumdiff/sumorig) could be greater than 1 and sim would be negative -- in those cases, the adjacent curves were clearly in different regional groups. Using sim < 0.7 to separate regions was relaxed for a few curves; these are identified with an asterisk in Table 1, which is located in the Supplementary Information. Transitional curves, with sim slightly less than 0.7, are plotted with dashed lines. To our knowledge the “sim” metric is not a standard metric. It is based on the calculation for finding the area between two data sets (the sumdiff term). Dividing by the sumorig term makes the output dimensionless. The metric was useful for color-coding the responses into different regions for illustration. The sim calculation was performed on the 70 dB data so the full frequency range could be used in the calculation.

We report from four gerbil cochleae, with a fifth in Supplemental Information. These preparations are used because responses were out of the noise at many pixels along an A-scan. The RL, BM, and OHC/DC-region responses observed here are consistent with those in previous studies from the gerbil hook region (22–24). What is new here is the observation and analysis of transitions between regions. Animal numbers are for laboratory record-keeping purposes.

## III. Results

### A. Introductory observations

Fig. 2 shows displacements from the BM, RL region, and OHC/DC region from G1047. The OHC/DC-region and RL-region responses are plotted along with the BM responses to facilitate comparisons. The OHC/DC region (Fig. 2A-C) initially is in-phase with the BM at the lowest measured frequencies, then a phase delay increases with increasing frequency (Fig. 2C). At 80 dB SPL, the phase of the OHC/DC region often shifts up to become in phase with the BM at higher frequencies, and a notch appears in the response magnitude, seen in Fig. 2B&C at ∼37 kHz. At ∼46 kHz, the phase shift and notch are even seen at 70 dB SPL. This behavior has been attributed to cancellation between BM and internal OHC motion, and subsequent dominance of BM motion (22). The OHC/DC-region gain displays sub-BF nonlinearity and boosting (increased responses relative to the BM responses). Sub-BF, the OHC/DC region has a larger magnitude response than the BM at all SPLs (Fig. 2B). In the BF peak, at 50 dB SPL, the OHC/DC region has a larger amplitude than the BM, but at higher SPLs, the OHC/DC region has a low-pass character without a notable peak. The RL region (Fig. 2D-F) has the highest BF responses, higher than those at the BM at all SPLs. Sub-BF, the RL responses are below BM responses. The RL phase leads the BM phase slightly throughout the frequency range. As has been observed in previous studies (23, 24), at frequencies below ∼20 kHz, the RL phase-lead re: BM was not present in all preparations; in-phase and slightly lagging phase were sometimes observed sub-BF and will be discussed below (Fig. 15).

**Fig. 2:**
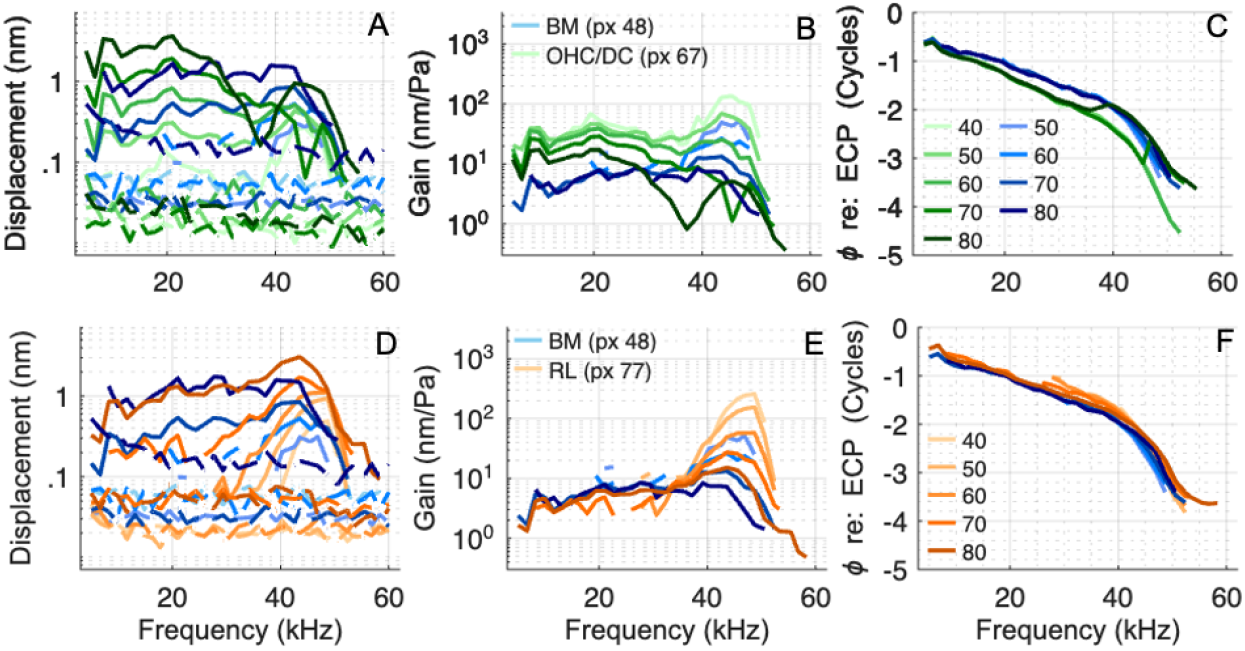
(A&D) BM (blue), OHC/DC-region (green), and RL-region (orange) displacement. (B&E) BM (blue), OHC/DC-region (green), and RL-region (orange) gain (displacement re: EC pressure (ECP)). The OHC/DC-region and RL-region gain is reported from 40-80 dB SPL, and the BM gain is reported from 50-80 dB SPL. (C&F) BM, OHC/DC-region, and RL-region phase re: ECP. Gerbil 1047 Run 5. BF = 45 kHz. Associated B-scan is in Fig. 3.

It is apparent from the regional displacements in Fig. 2 that significant strains will exist in the transitions between the RL, OHC/DC and BM regions of the OCC. In the remaining figures, we present displacement and strain results from four preparations, G1047, G1025, G1041, and G986 (and G1049 in Supplemental Information). Table 1 contains strain values at BF and BF/2. We begin with detailed sets of results from preparations G1047 and G1025.

### B. G1047

Fig. 3 shows a set of tuning curves from G1047, with BF ∼45 kHz. The *l,r,t* values were −0.62, −0.17, and 0.77, thus vertical distances in the B-scan are elongated from what they would be with a purely transverse view by a factor of ∼1.3 (figured using trigonometry as 1/[cos(arctan(0.62/0.77))], with the small radial component neglected in the calculation.) Panels A&B (pixels 49 and 58) are in the BM region, which also includes pixels 47 and 48 (consulting Table 1 and/or Fig. 5, which includes additional pixels). This is a 32 μm extent (figured as 11pixels x 2.7μm/pixel), corresponding to a 25 μm transverse extent (dividing 32 by 1.3 to account for the elongation). Panels C-F, pixels 63 to 73, are in the OHC/DC region of responses, which also includes pixel 62 (consulting Table 1 and/or Fig. 5). The spanned distance in the A-scan is 32 μm (corresponding to 25 μm transverse). Panel H (pixel 77), and pixel 78 are RL-region responses, and G (pixel 75) is termed intra-RL: the BF peak amplitude is like pixel 77, but sub-BF the amplitude is boosted and nonlinear, and the phase responses show increasing delay re: BM; these are characteristics like pixel 73 of the OHC/DC region (Fig. 3F).

**Fig. 3:**
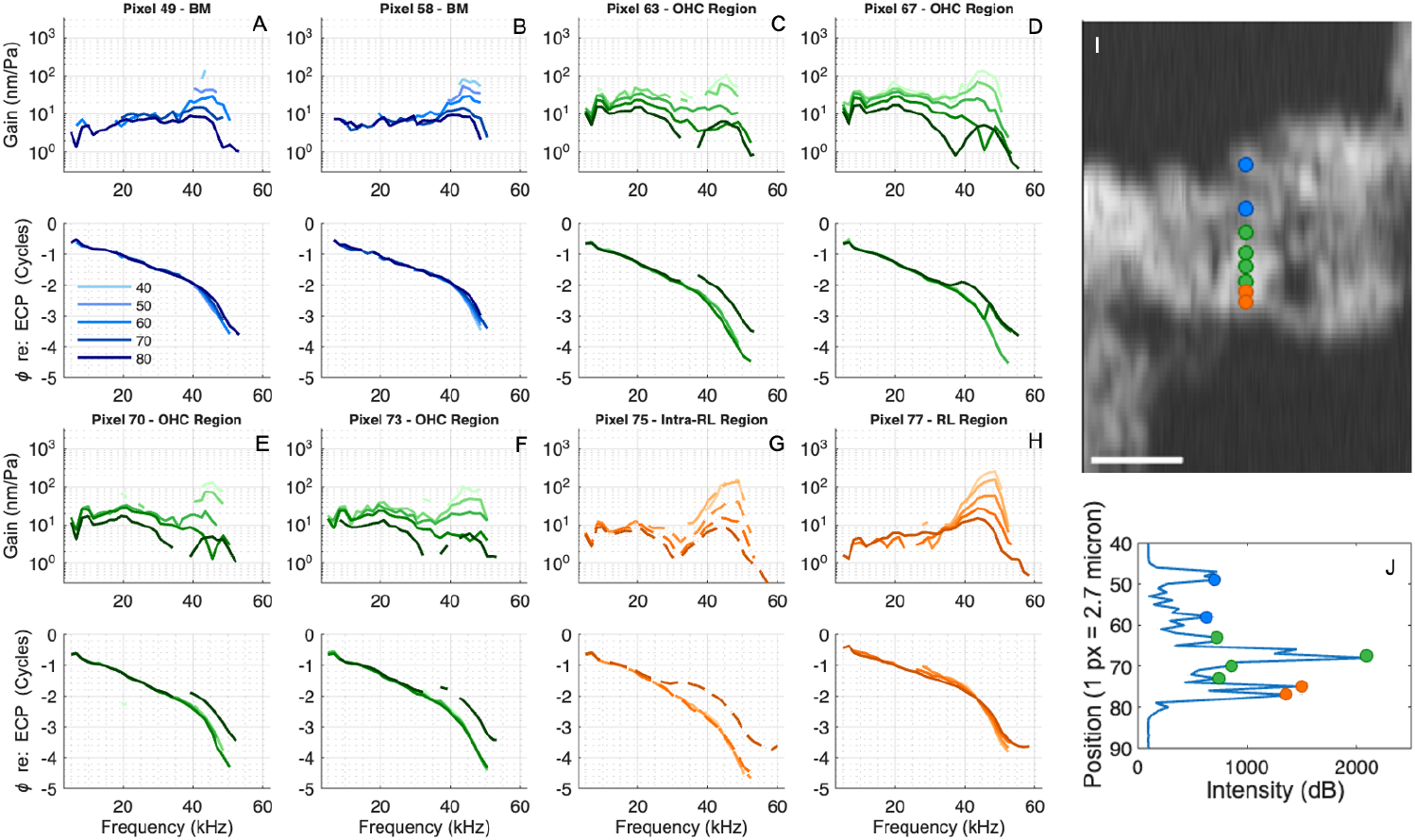
Responses from eight locations along a single A-scan (A-H). Gain responses are color-coded as follows: BM (blue), OHC-region (green), intra-RL-region (dashed orange), and RL-region (orange). The respective phase responses re: ECP are plotted below the gains. (I) B-scan with BM, OHC-region, intra-RL-region, and RL-region locations of measurements in (A-H) denoted with colored markers. Scale bar = 50 μm. (J) A-scan from 60 dB response (A-scans varied slightly during the course of a several-SPL run so we specify the SPL of the example A-scan). Optical axis components were (*l, r, t*) = (-0.62, -0.17, 0.77). Gerbil 1047 Run 5. BF = 45 kHz.

From these observations, steep yet graded variations occur between pixels 77, 75, and 73 in the RL and OHC/DC regions - an inclusive distance of 13.5 μm (10 μm transverse). The OHC/DC and BM regions both have relatively large spans of quite uniform motion. From pixel 78 (at the RL surface) to pixel 62 (the OHC/DC-region point closest to the BM region) is a distance of 46 μm (35 μm transverse). However, the OHC length at this BF location is expected to be only ∼20 μm (30); this underscores the need for the designation “OHC/DC” as a region that extends beyond the OHC’s basal end.

Fig. 4 shows a subset of the data in SPL-based groups at 50 and 70 dB SPL. These two SPLs are emphasized because 70 dB has many full-frequency data sets but still has features of activity that are less observable at 80 dB; 50 dB responses are useful to observe BF-peak responses at a low-moderate SPL. The purpose of this figure is to clearly show regional differences in the frequency responses, and to illustrate how the transition between regions differs at the two SPLs. Contrary to 70 dB, 50 dB OHC/DC-region responses retained a BF peak (Fig. 4A&C). At 50 dB, the RL-region peak and phase responses were nearly unchanged from pixels 77 to 75 (Fig. 4C&D), whereas at 70 dB SPL, pixel 75 has the “intra-RL” character described above (Fig. 4A&B). Thus, the “RL region” was wider at 50 than at 70 dB SPL (pixels 78 through 75 at 50 dB, pixels 78 and 77 at 70 dB).

**Fig. 4:**
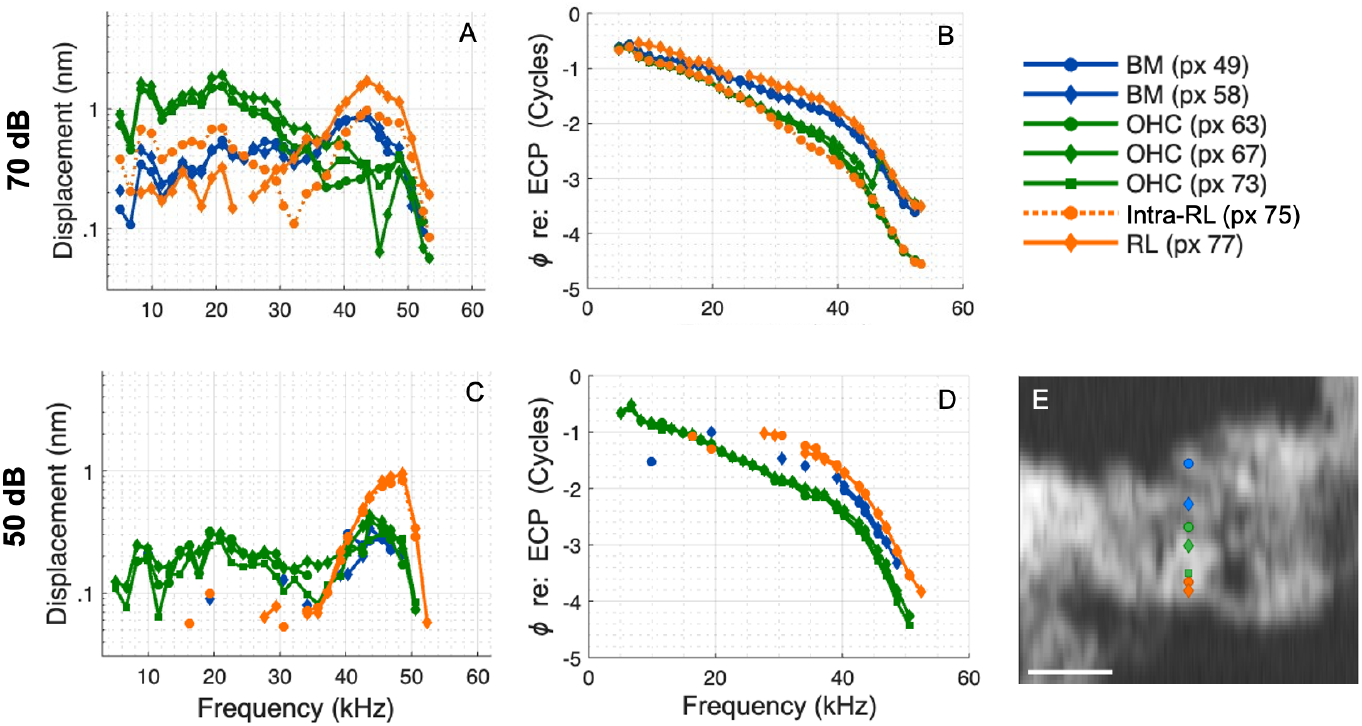
Displacement frequency responses at 50 and 70 dB from seven locations along a single A-scan. (A&C) BM (blue), OHC-region (green), intra-RL-region (dashed orange), and RL-region (orange) displacements at 70 dB SPL (A) and 50 dB SPL (C). (B&D) The respective phase responses re: ECP. (E) B-scan with BM, OHC-region, intra-RL-region, and RL-region locations of measurements reported in (A-D) identified. Scale bar = 50 μm. Gerbil 1047 Run 5. BF = 45 kHz.

Fig. 5 shows 1D heat-maps of: (column 1) displacement phase re: BM, with values between 0 and -1 cycle, (column 2) displacement amplitude, and (column 3) strain amplitude. The top two rows are 70 dB data at BF and BF/2 respectively (Fig. 5A-F), and the bottom row is 50 dB data at BF (Fig. 5G-I). The strain values are calculated from displacement values separated by 1 to 3 pixels. If the distance between available pixels was greater than 3, the strain value is considered an average strain, and these cases are delineated by a white frame, as in panel I. When adjacent displacement values were close to the noise level and/or almost identical in magnitude and phase, the resulting strain was sometimes in the noise (as determined by randomness in the strain phase) and excluded from the strain plot. After considering the displacement values, these gaps in the strain plots can often be taken as very low strain, even though numerical values (and thus colors) are missing. For numerical displacement values, Figs. 3 and 4 are most useful. The heatmaps of Fig. 5 are most usefully observed for regions of similar and different displacement and strain, coded by color, and we describe them in these terms. The significance of the numerical strain value is considered in the Discussion.

**Fig. 5:**
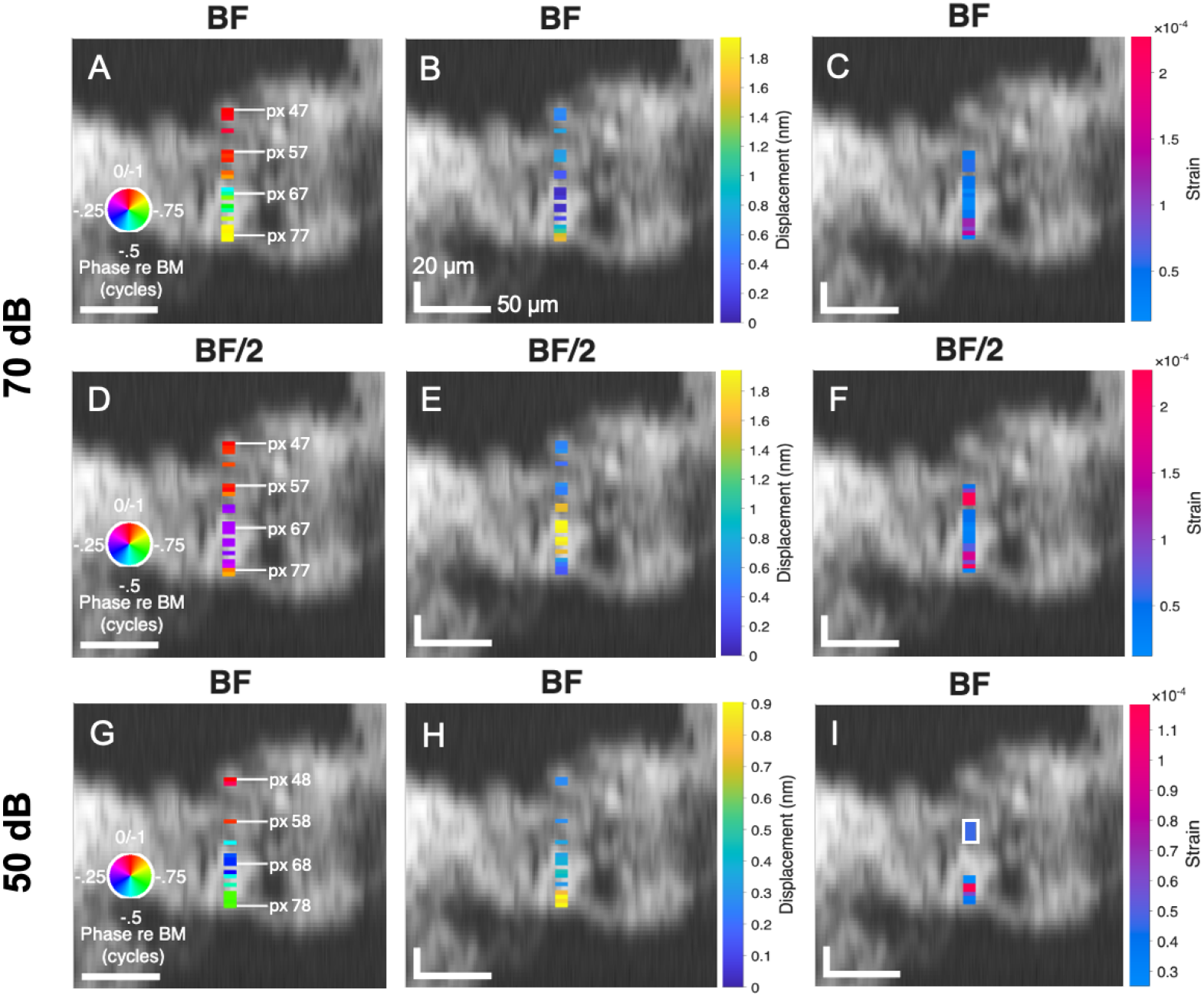
Displacement amplitude and phase and strain values along one A-scan, plotted as a 1D heat map onto the B-scan. (A, D, G) Displacement phase re BM: BF phase re: BM at 70 (A) and 50 dB (G); (D) BF/2 phase re: BM at 70 dB. (A,D used pixel 47, G used pixel 48 for BM reference.) (B, E, H) Displacement at BF at 70 (B) and 50 dB (H), and at BF/2 at 70 dB (E). (C, F, I) Displacement strain at BF at 70 (C) and 50 dB (I), and at BF/2 at 70 dB (F). The white rectangle in (I) denotes a region where the strain was calculated over more than 3 pixels. Vertical scale bars = 20 μm and horizontal scale bars = 50 μm. Gerbil 1047 Run 5. BF = 45 kHz.

In all rows, the BM region is clear in the relatively extensive region of bright red phases in column 1, coupled to a similar extent of mid-blue displacement amplitude in column 2. Many of the strain values in the BM region were in the noise - the BM region moved together, with little displacement strain.

At 70 dB and BF/2 (middle row), an extensive region of purple phases deeper than the red phases of the BM indicates a ∼0.23 cycle delay of the OHC/DC region (Fig. 5D). This is aligned with an extensive region of yellow (high) displacement amplitude in column 2 (Fig. 5E). This is aligned, in column 3, to cobalt blue, indicating relatively low strain -- the OHC/DC region appears to be moving almost as a body (Fig. 5F). Close to the RL, a two-pixel-wide (78 and 77) region of orange phase (Fig. 5D) indicating a ∼0.9 cycle lag (better interpreted as a 0.1 cycle lead) is coupled to blue (low) amplitude values in column 2 (Fig. 5E); adjacent to these (shallower pixels 76 and 75) the phase changes to a purple lag (Fig. 5D) and the displacement amplitude goes briefly to blue-green (pixel 75), then to the yellow (higher) values of the OHC/DC region (Fig. 5E). Through this RL-to-OHC/DC transition, the strain has two blue bands (low strain) separated by two magenta bands (high strain) (Fig. 5F). The deepest blue strain band, from the difference of pixels 78 and 77, indicates that these two positions closest to the RL boundary moved together - a distance of ∼5 μm (4 μm transverse). The deepest magenta high-strain band occurs due to changes in both the phase and amplitude of the displacement, the second (shallower) high-strain band is due to a spatial variation in displacement amplitude. A third (shallowest) magenta high-strain band, around pixel 60, appears at the transition between the large sub-BF OHC-region displacement and small sub-BF BM displacement (Fig. 5E&F).

At 70 dB and BF (top row), deeper than the BM, the green displacement phase indicates a ∼0.75 cycle delay of the OHC/DC region relative to the BM (Fig. 5A). The displacement amplitude in this region codes dark blue; displacement is smaller than at the BM (Fig. 5B). With increasing depth the blue-green-yellow shift in displacement amplitude shows the transition to the high displacement amplitude of the RL. The yellow phase in this region (Fig. 5A), upon consideration of Fig. 4B, is best considered a small phase lead re: BM, not a large lag. Close to the RL, the strain is high in two dark magenta bands (Fig. 5C), similar to the bands at BF/2 (Fig. 5F). Shallower than these magenta bands, the strain remains low (blue) (Fig. 5C) - there is not a third magenta band around pixel 60 as there was at BF/2 (Fig. 5F); the transition between OHC/DC and BM was evident in Fig. 4A, but the displacement values are relatively small and phase difference approaching a full cycle, leading to relatively small strain (Fig. 5C).

At 50 dB and BF (bottom row), the displacement phase transitions red/blue/green with increasing depth (Fig. 5G). Here, the blue indicates OHC/DC-region phase is delayed from the BM by only ∼0.35 cycle, less than it was delayed at BF and 70 dB (consult Fig. 4, Fig. 5A). At 50 dB, the OHC/DC region retains a prominent BF peak, slightly larger than the BM peak (Fig. 4C) – color-wise, the OHC/DC displacement values are blue-green, versus the cobalt-blue of the BM (Fig. 5H). The blue phase (Fig. 5G) and blue-green displacement of the OHC/DC region (Fig. 5H) span pixels 66 to 70 – there was little spatial variation in displacement and the spatial differences were in the noise, accounting for the gap in the strain map (Fig. 5I). The RL-region displacement was relatively high (yellow) (Fig. 5H) and the green phase indicates a lead re: BM of ∼0.3 cycle over pixels 78 to 75, about 11 μm (8 μm transverse) (Fig. 5G). A band of high (magenta) strain appears between pixels 73 and 75 (Fig. 5I).

Strain has both amplitude and phase, and Fig. 6 shows strain frequency responses found from displacement at pixels close to the RL. 70 dB data are shown, with displacement in A&B and strain in C&D. The strain amplitude-vs-frequency (Fig. 6C) has a gentle bimodal shape with the peak in the BF region due to the transition between large RL and small OHC/DC displacement amplitudes (Fig. 6A), and a peak at frequencies around 20 kHz due to the transition between small RL and large OHC/DC displacement amplitudes. The strain phase (Fig. 6D) is similar for the three calculated strains. The strain phase 77-76, when copied to panel B, aligns with the displacement phase of pixel 77 (close to the RL surface) from 25 to 50 kHz. Below ∼ 20 kHz, the strain leads the displacement of pixel 77 by ∼0.2 cycle and is ∼1/2 cycle off from the phase of displacement pixels 73, 75, 76.

**Fig. 6:**
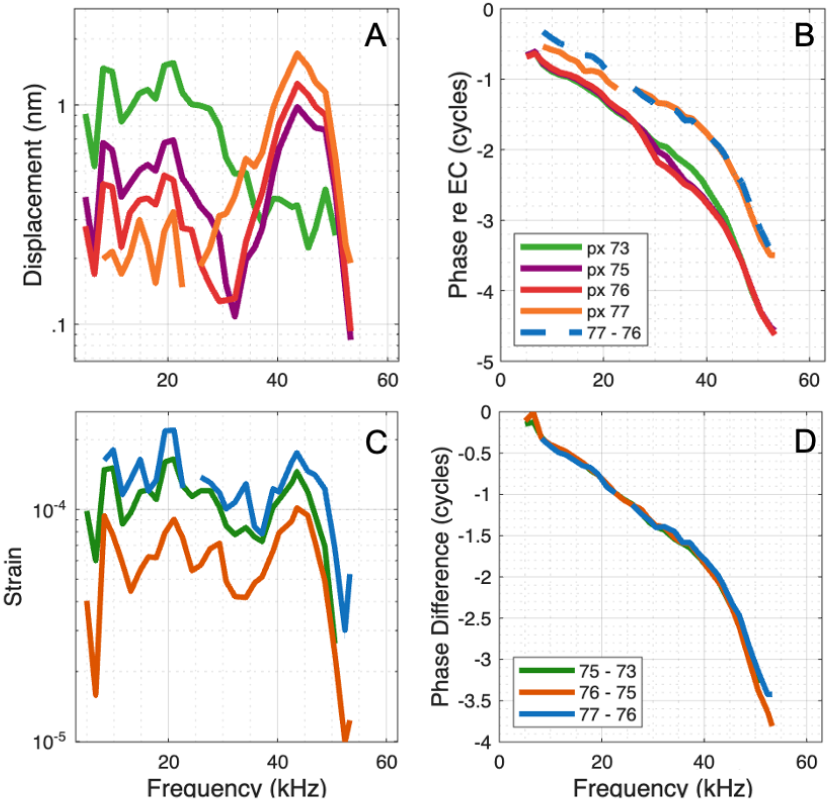
Strain and displacement responses close to the RL at 70 dB. (A&B) The displacement amplitude and phase re: ECP at four pixels close to the RL. (C&D) The strain amplitude and strain phase calculated between the pixels in (A&B). The strain phase found with pixels 77 and 76 is plotted in (B) as a blue dashed line for comparison. Gerbil 1047 Run 5. BF = 45 kHz.

### C. G1025

The BF was ∼45.5 kHz and *l,r,t* values were -0.64, -0.1, and 0.76, and similar to G1047, the B-scan is elongated from what it would be with a purely transverse view by a factor of ∼1.3. Fig. 7 shows a set of gain tuning curves from G1025, and Fig. 8 shows displacement tuning curves sorted to 50 and 70 dB SPL.

**Fig. 7:**
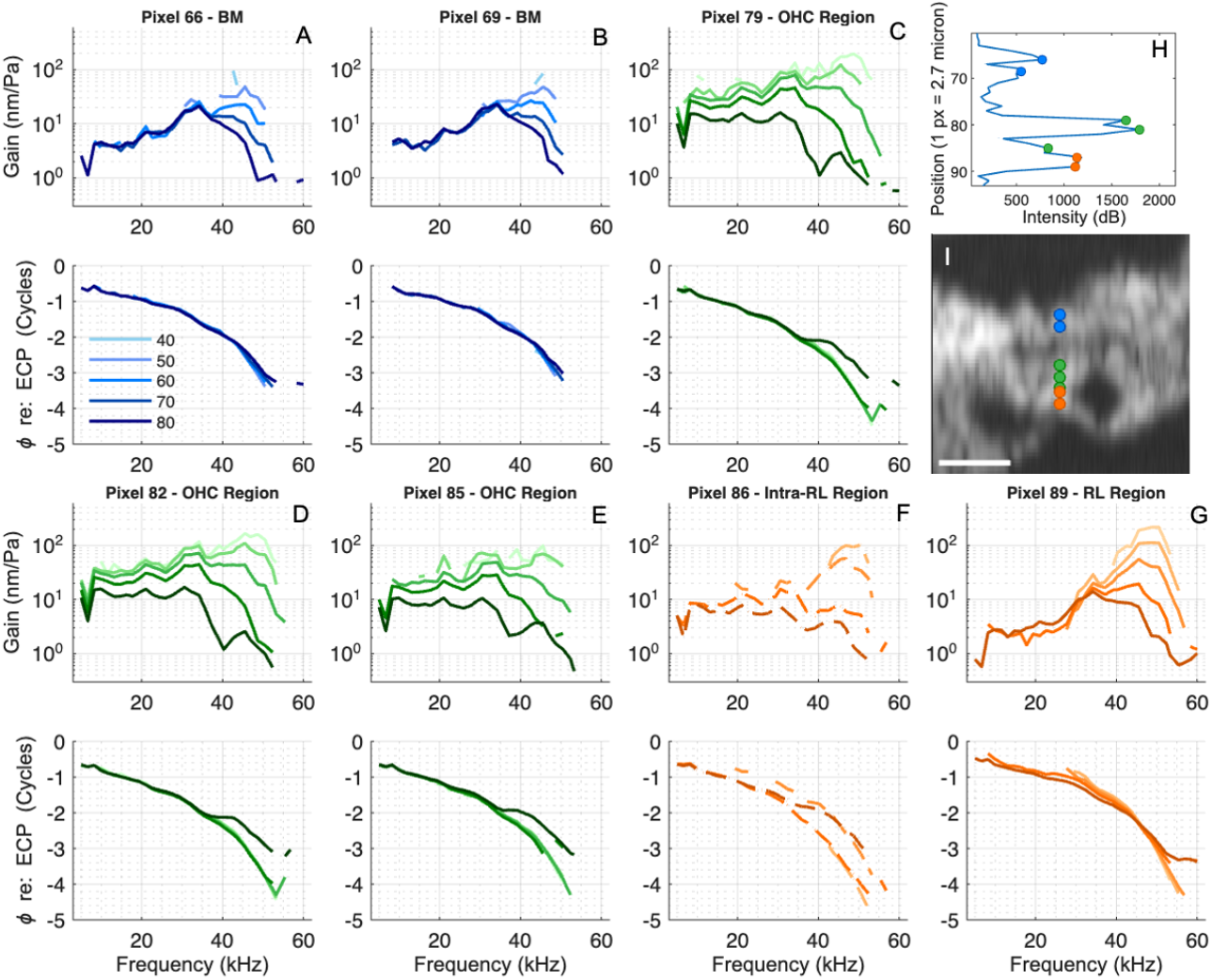
Responses from seven locations along a single A-scan. (A-G) Gain responses color-coded as: BM (blue), OHC-region (green), intra-RL-region (dashed orange), and RL-region (orange). The respective phase responses re: ECP are plotted below the gain responses. (H) A-scan of 70 dB response. (I) B-scan with BM, OHC-region, intra-RL-region, and RL-region locations of measurements reported in (A-G) denoted with colored markers. Scale bar = 50 μm. Optical axis components were (*l, r, t*) = (-.64, -0.1, 0.76). Gerbil 1025 Run 13. BF = 45.5 kHz.

**Fig. 8:**
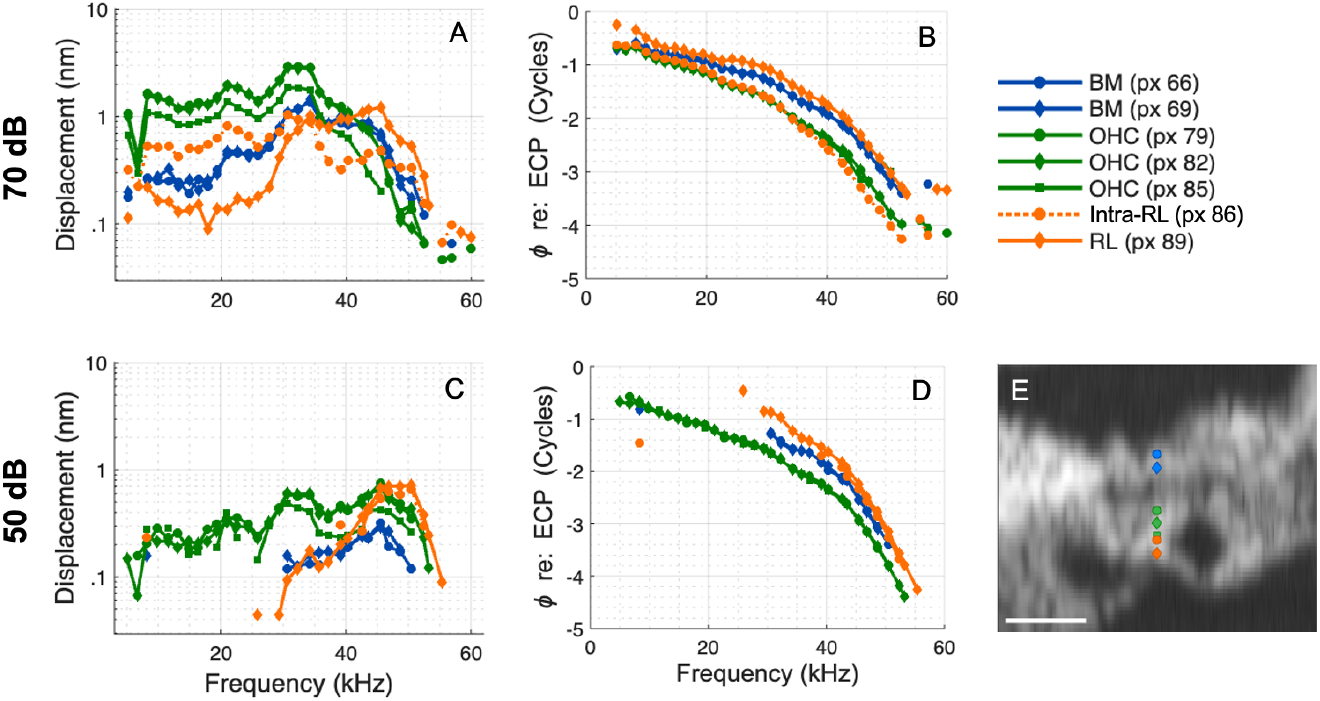
Displacement frequency responses at 50 and 70 dB from seven locations along a single A-scan. Caption as in Fig. 5. Gerbil 1025 Run 13. BF = 45.5 kHz.

Pixel 89 is close to the RL surface and 86, 3 pixels shallower, appears as an “intra-RL” point -- at 70 dB retaining a small BF peak but also showing OHC/DC-like sub-BF boosting (Fig. 8A). Pixel 85 has the form of the OHC/DC-region pixels 82 and 79 (Fig. 8A), but at a reduced amplitude. Considering Fig. 8, at 50 dB, pixel 86 is more closely aligned with the RL than it was at 70 dB; a similar observation was made in Fig. 4. The heatmaps in Fig. 9 echo those from G1047 (Fig. 4), with minor color shifts. The wide purple OHC/DC-region phases at 70 dB, BF/2 in Fig. 5D are slightly bluer in Fig. 9D, indicating a phase lag re: BM of ∼0.25 cycle. In panel H (50 dB displacement amplitude), a green-yellow color of relatively high displacement at pixel 86 was obscured against the gray B-scan so the region is enlarged in an inset. This panel (H) seems to differ most between G1025 and G1047, with the region of high (yellow) displacement extending from the RL edge through the OHC/DC region in G1025. Comparing Figs. 8 and 4, the OHC/DC 50 dB BF displacement amplitude in G1025 (Fig. 8C) is close to that of the RL, whereas in G1047 (Fig. 4C) it was lower, close to that of the BM. Despite the similar RL and OHC/DC displacement amplitudes in Fig. 8H, the strain in panel I shows a magenta band, due to the substantial phase change between “intra-RL” (pixel 86) and OHC/DC (pixel 85) (green, ∼0.25 cycle lead re: BM at pixel 86 and blue, ∼0.4 cycle lag at pixel 85). At 70 dB, the region of strain close to the RL is similar at BF and BF/2 (Fig. 9C&F) and wider than at 50 dB (Fig. 9I), observations that are similar to G1047. At BF/2, there is a region of low strain from pixels 82 to 78 (Fig. 9F). This is within the OHC/DC region, with yellow displacement amplitude (Fig. 9E) and purple displacement phase (Fig. 9D). Pixel 78, the shallowest pixel in the OHC/DC region, is 30 μm (23 μm transverse) from pixel 89 close to the RL surface - as noted above, this distance is larger than the expected ∼ 20 μm OHC length, so includes part of the DC. At BF/2, a region where the average strain is relatively high (purple) is set off in a box from pixel 78 to 71 (due to lacking intermediate data) (Fig. 9F), where high sub-BF OHC/DC-region displacement transitions to BM-region displacement. At BF and 70 dB, the strain is relatively low at pixels shallower than 82 (Fig. 9C).

**Fig. 9:**
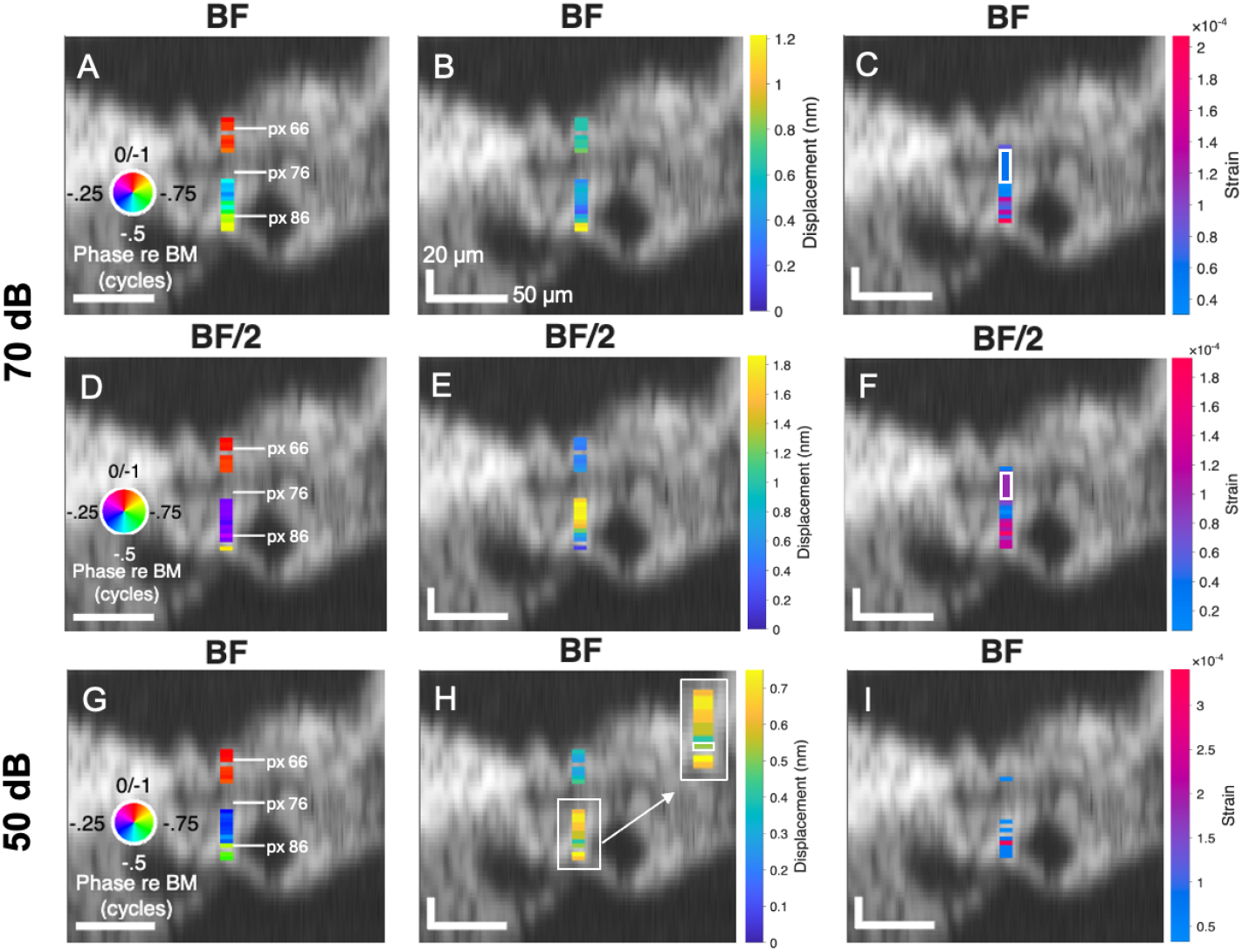
Displacement amplitude and phase at all pixels deemed out of the noise along one A-scan, plotted as a 1D heat map onto the B-scan. (A, D, G) Displacement phase re: BM: BF phase re: BM at 70 (A) and 50 dB (G); (D) BF/2 phase re: BM at 70 dB. (Pixel 64 used for BM reference.) (B, E, H) Displacement at BF at 70 (B) and 50 dB (H), and at BF/2 at 70 dB (E). (C, F, I) Displacement strain at BF at 70 (C) and 50 dB (I), and at BF/2 at 70 dB (F). Gerbil 1025 Run 13. BF = 45.5 kHz. A strain frequency response plot for G1025 (like that of Fig.6) is included in the Supplemental Information.

### D. G1041

Figs. 10-12 show frequency response curves and heat maps for G1041, with BF 43.6 kHz. *l,r,t* values were −0.57, −0.39, 0.72, so vertical distances in the B-scan are elongated from what they would be with a purely transverse view by a factor of ∼1.4. Overall, the trends noted from G1047 and G1025 above are maintained. Pixel 88 appears to be deeper than the main structure of the RL (Fig. 10I) and might be the edge of the TM. From Fig. 11B&D, at both 70 and 50 dB, the RL-region phase-lead re: BM is slightly smaller than in G1047 (Fig. 4B&D) and G1025 (Fig. 8B&D).

**Fig. 10:**
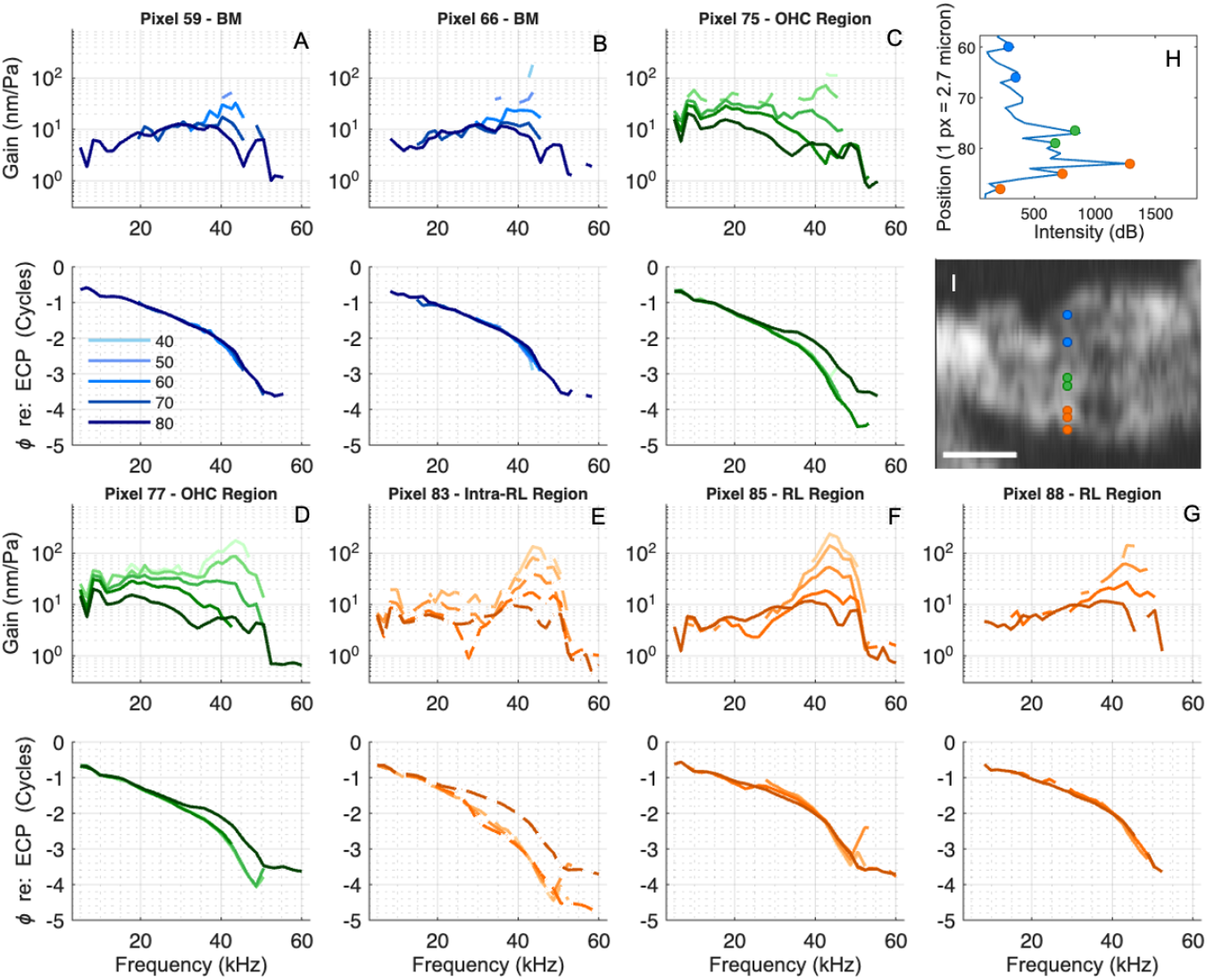
Responses from seven locations along a single A-scan. (A-G) BM (blue), intra-OHC-region (dashed green), OHC-region (green), intra-RL-region (dashed orange), and RL-region (orange) gain. The respective phase responses of (A-G) re: ECP are plotted below the gain responses. (H) A-scan of 50 dB response. (I) B-scan with BM, intra-OHC-region, OHC-region, intra-RL-region, and RL-region locations of measurements reported in (A-G) are denoted with colored markers. Scale bar = 50 μm. Optical axis components were (*l, r, t*) = (−0.57, −0.39, 0.72). Gerbil 1041 Run 6. BF = 43.6 kHz.

**Fig. 11:**
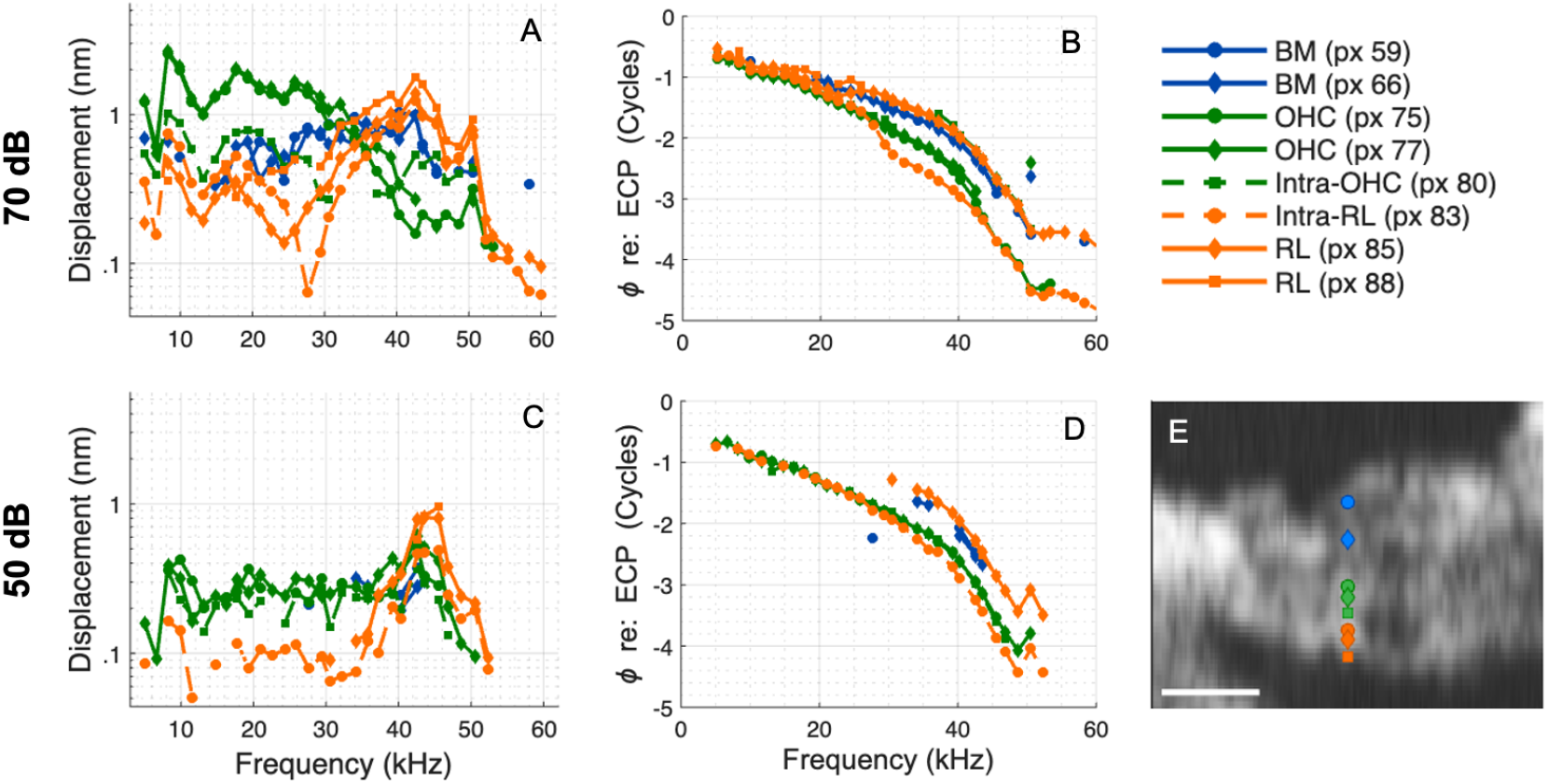
Displacement frequency responses at 50 and 70 dB from seven locations along a single A-scan. Caption as in Fig. 5. Gerbil 1041 Run 6. BF = 43.6 kHz.

In Figs. 10 and 11, the transition from RL to OHC/DC proceeds through pixels 88, 85, 83, 80, and 77. Pixel 88 displays characteristic RL motion, with a high magnitude at the BF peak and generally a small phase lead re: BM. Pixel 85 is similar to 88 at 50 dB (Fig. 11C&D) and reasonably similar at 70 dB (Fig. 11A&B), but begins to show intra-RL behaviour, displaying an amplitude notch at ∼24 kHz (Figs. 10F&11A), below which the phase lags the BM slightly (Fig. 11B). Intra-RL pixel 83 has notch and phase transition at around 28 kHz (Figs. 10E&11A), and compared to pixels 85 and 88, 83 has a greater sub-BF amplitude and/or a slightly lower response at the BF peak at 50 and 70 dB (Fig. 11A&C). In pixel 83, the phase transitions to greater delay compared to pixel 85, and at frequencies near BF is a full cycle beneath the pixel 85 phase (thus, these two pixels are approximately in phase at BF) (Fig. 11B&D). At frequencies below the notch, pixel 83 phase lags BM (Fig.11B). Pixel 80 is another intermediate position, with sub-BF responses smaller than OHC/DC responses (for example, pixel 77), but greater than the intra-RL responses at pixel 83 (Fig. 11A). At 70 dB, pixel 80 has a more prominent BF peak than OHC/DC-region pixel 77 (Fig. 11A), but the peak is smaller than the intra-RL and RL regions at pixels 83 and 85. Pixel 80 phase is like the RL region in the BF peak and like the OHC/DC sub-BF (70 dB) (Fig. 11B).

Considering the 1D heatmaps of Fig. 12 at 70 dB (top two rows), pixels 85 to 80 are part of the transitional region. At BF, there are moderate (purple) strains close to the RL and over several pixels strain increases with distance from the RL (Fig. 12C). The largest (magenta) strain occurred between the “intra-OHC” pixel 80 and OHC/DC region at pixel 76 (BF, Fig. 12C) or 77 (BF/2, Fig. 12F). Between RL-region pixels 88 and 85, at 70 dB there is little change in displacement phase (Fig. 12A&D) but a small reduction in amplitude (Fig. 12B&E), leading to moderate (purple) strain at BF (Fig. 12C) and smaller (blue) strain at BF/2 (Fig. 12F). At 50 dB and BF (bottom row), between pixels 88 and 85 there was little change in amplitude or phase (Fig. 12G&H), leading to a small strain value that was indiscernible from noise, thus it was not color-coded (Fig. 12I). A relatively large (magenta) strain exists between pixels 85 and 84.

**Fig. 12:**
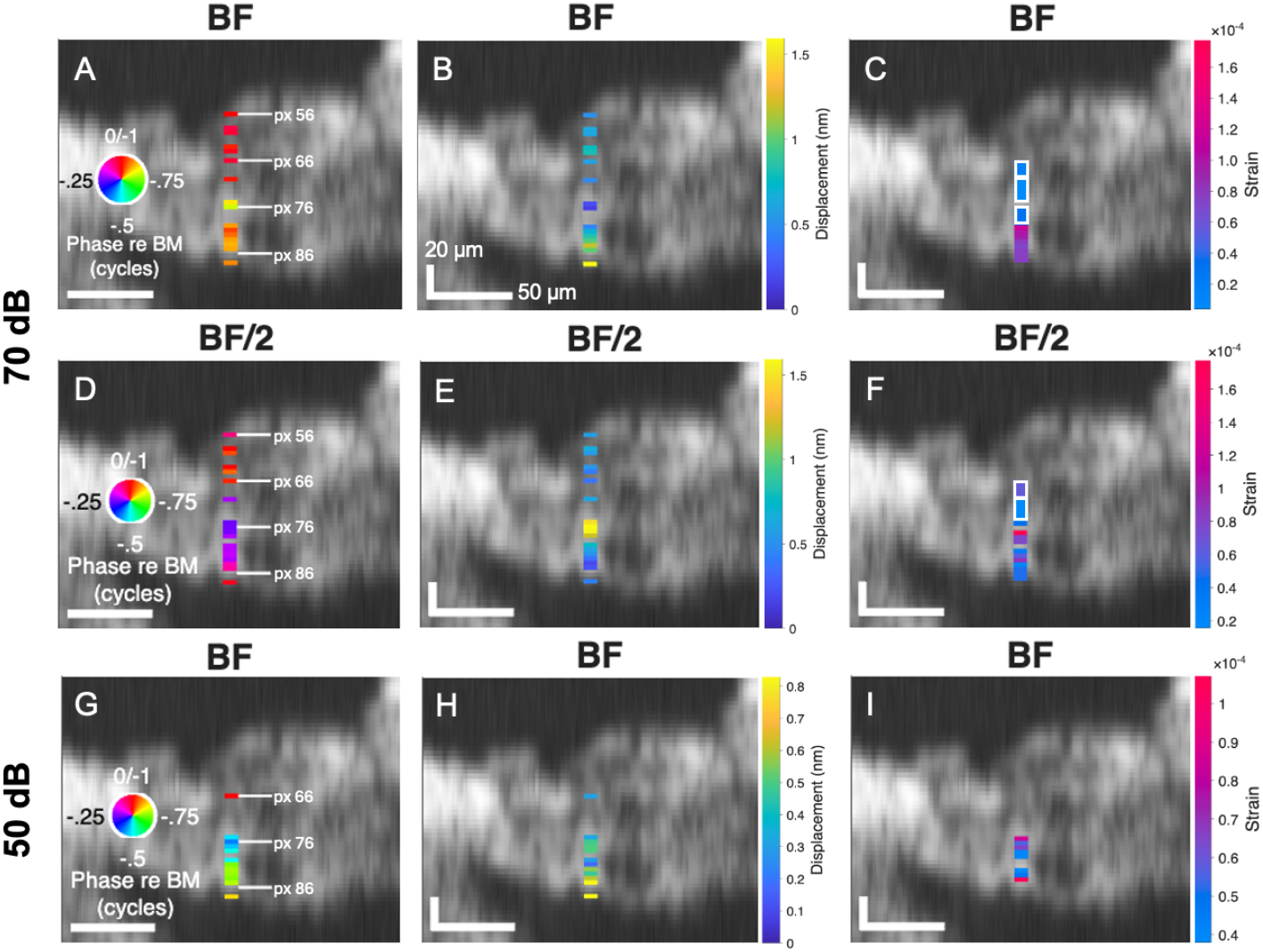
Displacement amplitude and phase at all pixels deemed out of the noise along one A-scan, plotted as a 1D heat map onto the B-scan. (A, D, G) Displacement phase re BM: BF phase re: BM at 70 (A) and 50 dB (G); (D) BF/2 phase re: BM at 70 dB. (A,D used pixel 59, G used pixel 66 for BM reference.) (B, E, H) Displacement at BF at 70 (B) and 50 dB (H), and at BF/2 at 70 dB (E). (C, F, I) Displacement strain at BF at 70 (C) and 50 dB (I), and at BF/2 at 70 dB (F). Gerbil 1041 Run 6. BF = 43.6 kHz.

### E. G986

Figs. 13&14 are the final group of data figures, from G986, with BF 44.5. *l,r,t* values were -0.48, -0.35, 0.8, thus vertical distances in the B-scan are elongated from what they would be with a purely transverse view by a factor of ∼1.25. This experiment was performed as part of a previous set, in which we selected a series of local maxima in the A-scan as the pixels to be analyzed for displacement. This results in a relatively sparse data set, which was nevertheless a useful, supportive set. (Some of these data were presented in a different format in Strimbu et al., 2024.) The displacement frequency responses sorted to 50 and 70 dB are not included here, but can be found in the Supplementary Information. The trends observed above are apparent. For example, in the heatmaps of Fig. 14, at 70 dB and BF/2, there is a wide region of yellow (relatively high) displacement amplitude (Fig. 14E) and purple phase (∼0.23 cycle lag) (Fig. 14D), corresponding to the OHC/DC region (here spanning at least pixels 74 to 65, a distance of 30 μm (24 μm transverse). RL-region pixel 76 is slightly within the surface of the RL, and it has some “intra-RL” characteristics. For example, there is a notch in the 70 dB amplitude at ∼23 kHz (Fig. 13F); the phase leads the BM only above this frequency. There is a sudden transition between RL-region and OHC/DC-region behavior between pixels 76 and 74, giving rise to the bright magenta strain value in the third column of all rows (Fig. 14C,F,I). At BF, the only region of magenta strain is between pixels 76 and 74 (Fig. 14C&I); at BF/2 a second purple band appears between the BM and OHC/DC regions (Fig. 14F), with the transition from the large (yellow) OHC/DC displacement to the smaller (blue) displacement of the BM (Fig. 14E).

**Fig. 13:**
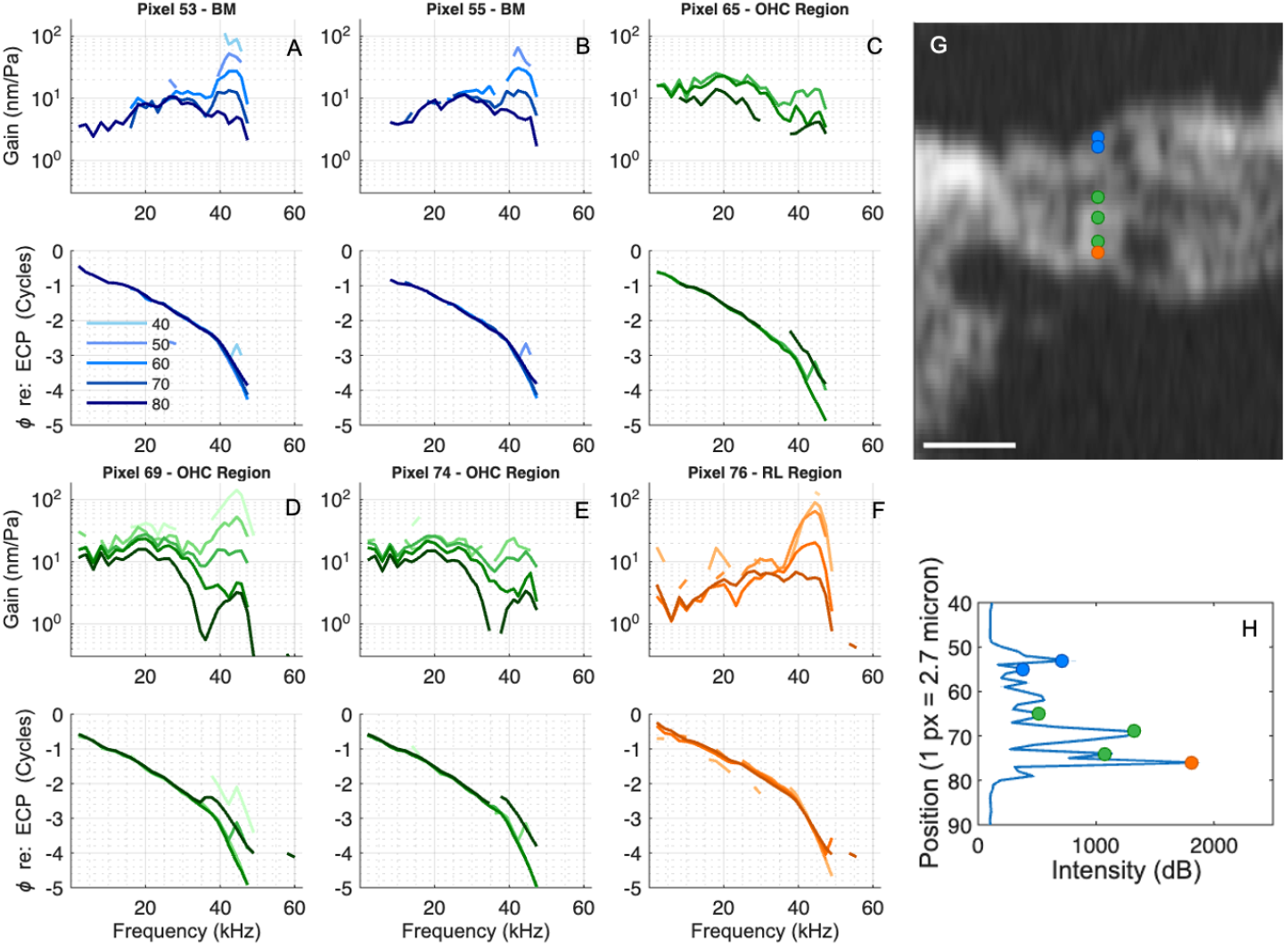
Responses from six locations along a single A-scan. (A-F) BM (blue), OHC-region (green), and RL-region (orange) gain. The respective phase responses of (A-F) re: ECP are plotted below the gain responses. (G) B-scan with BM, OHC-region, and RL-region locations of measurements reported in (A-F) are denoted with colored markers. Scale bar = 50 μm. (H) A-scan of 50 dB response. Optical axis components were (*l, r, t*) = (-0.48, -0.35, 0.8). Gerbil 986 Run 4. BF = 44.5 kHz.

**Fig. 14:**
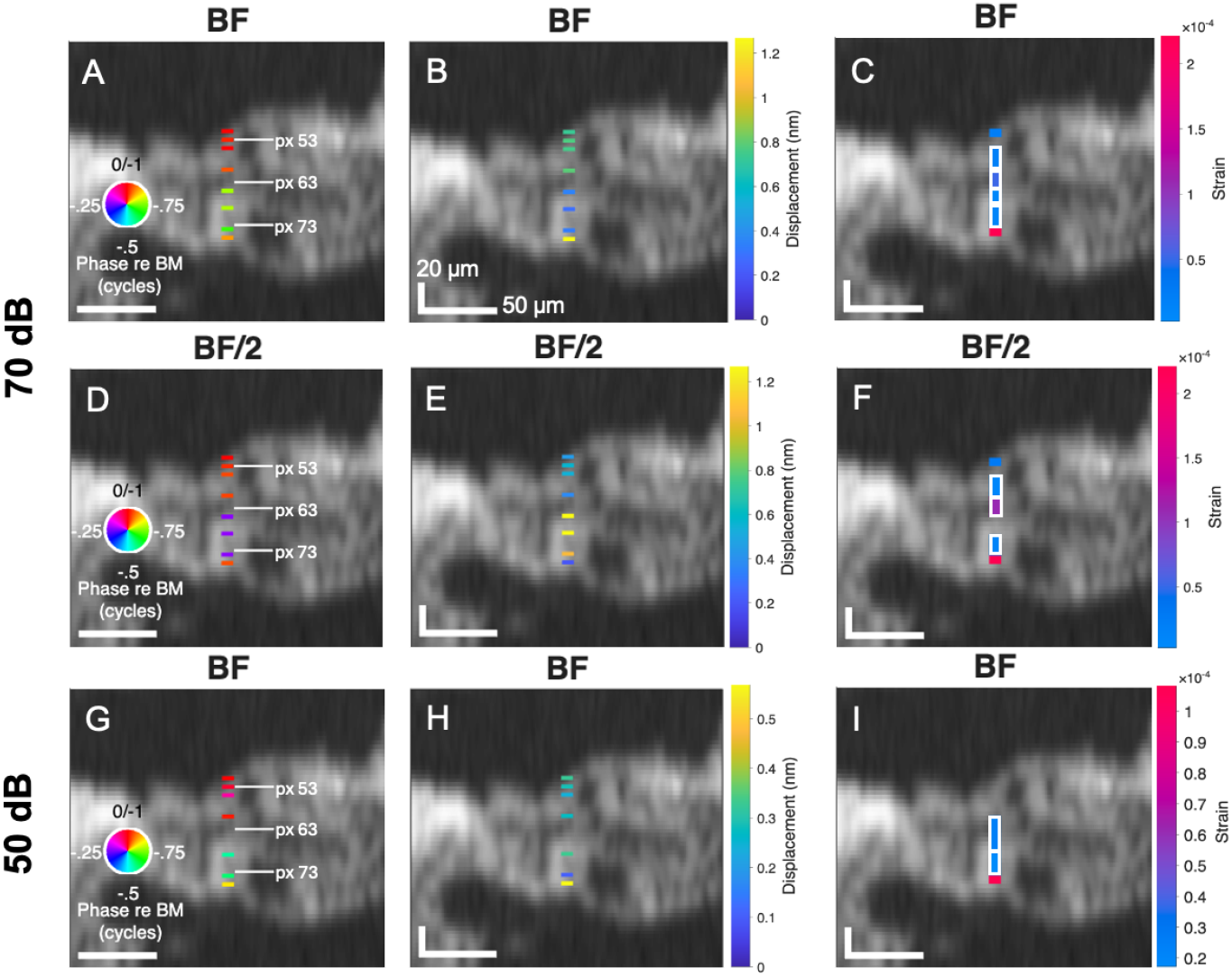
Displacement amplitude and phase at all pixels deemed out of the noise along one A-scan, plotted as a 1D heat map onto the B-scan. (A, D, G) Displacement phase re BM: BF phase re: BM at 70 (A) and 50 dB (G); (D) BF/2 phase re: BM at 70 dB. (Pixel 51 was used for BM reference.) (B, E, H) Displacement at BF at 70 (B) and 50 dB (H), and at BF/2 at 70 dB (E). (C, F, I) Displacement strain at BF at 70 (C) and 50 dB (I), and at BF/2 at 70 dB (F). Gerbil 986 Run 4. BF = 44.5 kHz.

## IV. Discussion

## A. Summary of observations related to OCC anatomy

The mechanics of the cochlea produce amplification of a sound-induced nanometer-scale fluid-mechanical wave, traveling from the base of the cochlea towards its apex. OCC motion, until recently observable only at the BM in the high frequency cochlear base, has now been observed deeply within the OCC. The RL is the deepest region of the OC, adjacent to the stereocilia, the site of mechanoelectric transduction. Unlike the OHC/DC region, in the base of the gerbil cochlea the RL displays little/no sub-BF boosting and the phase-frequency slope is relatively similar to that of the BM. The similarity of BM and RL motion is likely related to their connection via the structural, microtubule-rich supporting cells, PCs and DCs. The BM is a primary structural component of the OCC, but in point-stiffness measurements, the cells of the OC increased the stiffness measured at the BM by a factor of about two generally, and of about 10 at the outer PC (31–33). In measurements and analysis in excised gerbil cochleae, the DCs were found to have a stiffness ∼2.5 times that of the OHCs (34). The RL is a tiled structure formed by the apices of HCs, PCs, and the phalangeal processes of DCs. The apices of all these cells are rich in the structural protein actin. In HCs, the cuticular plate anchors the stereocilia and has been characterized as a dense actin gel (35). Actin gels can have significant stiffness, depending on the actin density and cross-linking molecules (8, 36), and the RL is considered to be stiff (34), although in recent observations, it was not completely rigid (24).

OHCs are the pumping heart of cochlear amplification. Voltage-induced stresses in the OHC lateral membrane produce electromotility: contraction and elongation along the OHC length. In excised OHCs, the electromotile amplitude for an AC stimulus up to 50 kHz was ∼1nm/mV for an OHC of length ∼50 µm, corresponding to a strain of ∼2*10^-5^ / mV (17, 19). The largest strains measured in the current study were generally ∼1*10^-4^ at 50 dB SPL, at the BF. To compare the excised OHC strains to in vivo strains, we estimate OHC transmembrane voltage at 50 dB SPL using results from the literature: Starting with extracellular voltage measured close to the BM in gerbil, OHC transmembrane voltage was estimated as ∼2 mV at ∼30-40 dB SPL at the 20 kHz BF (37). In in vivo intracellular measurements in guinea pig, OHC AC receptor potentials were ∼ 1 mV at 30 dB at the cell’s 1 kHz BF (38) and ∼ 5 mV at 50 dB at the cell’s 16 kHz BF (39). Based on those quantities, at 50 dB SPL we expect OHC transmembrane voltage at the BF to be ∼5 mV. Then our measured strains at 50 dB are ∼1*10^-4^ /5 mV, or 2*10^-5^/mV, which is in agreement with the excised OHC strain values. However, at 50 dB, the relatively high strains in vivo were present over short distances compared to the ∼20 µm length of an OHC (30). In vivo OHC-based force is mechanically loaded by surrounding cells and structures, and the character and size of electromotility must reflect those loads.

In vivo, voltage-induced OHC strains and stresses work within the structural framework of the OCC, overpowering forces dissipating the traveling wave to permit it to travel further and attain a greatly increased BF peak. In the observations here, the region of largest strain was close to the RL. At 70 dB, it could extend a length that would include much of the OHC body; this strain is likely due to active contraction and elongation of the OHC body. Beyond that (shallower), the OHC/DC region could move nearly as a uniform body. Between the OHC/DC and BM regions, especially sub-BF, there was a second region of large strain. Based on the distances, this occurred well beyond the OHC body and cannot be considered to be direct OHC electromotile strain. Two primary observations of this study are the substantial strain extending into the OHC lateral region from RL - likely the strain of electromotility; and the nearly uniform region of OHC/DC motion – likely driven by electromotility. Either or both of these motions could be critical for the operation of the cochlear amplifier. The large OHC-region motion has been observed and discussed in previous studies of the gerbil base (22, 25, 26, 40). Below, we will consider these two observations in turn.

### B. Transition from RL to OHC/DC

The motions in the RL and OHC/DC regions differ in both amplitude and phase. The transition between these regions was relatively abrupt at 50 dB compared to 70 dB. For example, in G1047 pixels 78 and 77 at/close-to the RL moved similarly at 70 dB (an inclusive distance of 5.4 μm, 4.2 μm transverse), and adjacent pixel 76 was an “intra-RL” point (Fig. 5D,E,F). At 50 dB, pixels 78 to 75 moved similarly, an inclusive distance of 10.8 μm, 8.3 μm transverse (Fig. 5G,H,I); the next measurable pixel, 73, was in the OHC/DC region. “RL-region” motion also extended through more pixels at 50 than at 70 dB in G1047 (Fig. 4), G1025 (Fig. 8), and G1041 (Fig. 11). (In G986, due to relatively sparse sampling, the extent of the RL-region motion could not be evaluated. In G1049 in the Supporting Information the observation was not supported because at both 70 and 50 dB SPL a transition from RL-like to intra-RL occurred between the same two pixels (Figs. S4,S5).)

At 70 dB, the transition between RL and OHC/DC motion could proceed relatively smoothly, with intra-region points having characteristics intermediate between RL and OHC/DC regions. In the following, we show that the motion of intermediate positions can be approximately predicted by assuming a frequency-independent progression, expressed in Eq. 3, where *x* is a complex value that contains both magnitude and phase, represented as real and imaginary parts for the calculation. α = 0 corresponds to motion at the RL, α = 1 corresponds to the approximately uniform OHC/DC motion, 0 < α < 1 is motion between the RL and OHC/DC regions.

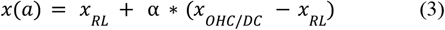

Fig. 15A&B show examples of the output of Eq. 3, using the RL-region and OHC/DC-region motions from G1047. Pixels 77 (which was more reflective, with cleaner data than pixel 78) and 67 were used for the RL and OHC/DC regions, respectively. (Supplemental Information contains examples using the RL and OHC/DC-region motions from G1041.) With α = 0.15 (Fig. 15A), a notch and phase re: BM cross-over occurs, similar to pixel 85 from G1041 at 70 dB (Fig. 15C). α = 0.45 generates a two-peaked response (Fig. 15B), with the intra-region phase beginning like OHC/DC and ending like RL (minus one full cycle). This behavior was observed at pixel 83 from G1041 (Fig. 11B), pixel 75 from G1047 (Fig. 15D), and pixel 86 from G1025 (Fig. 8B). In G1047, the RL surface was at pixel 78, and pixel 75 was ∼8 μm shallower (transverse distance, figured as 4 pixels x 2.7/1.3 μm). If the OHCs were expanding/contracting uniformly and were 20 μm long, we would expect α at this location to be ∼8/20 = 0.4, which is similar to 0.45 – so in this example α could be considered as a distance-based scaling parameter. Because of the large sub-BF motions of the OHC/DC region, even small α values can shift the low frequency phase from leading BM to slightly lagging, as observed in Fig. 15A, and this “entraining” effect of OHC/DC motion on RL motion could give rise to some of the variation in RL-region sub-BF phases observed in this study and in previous observations (22, 24).

**Fig. 15:**
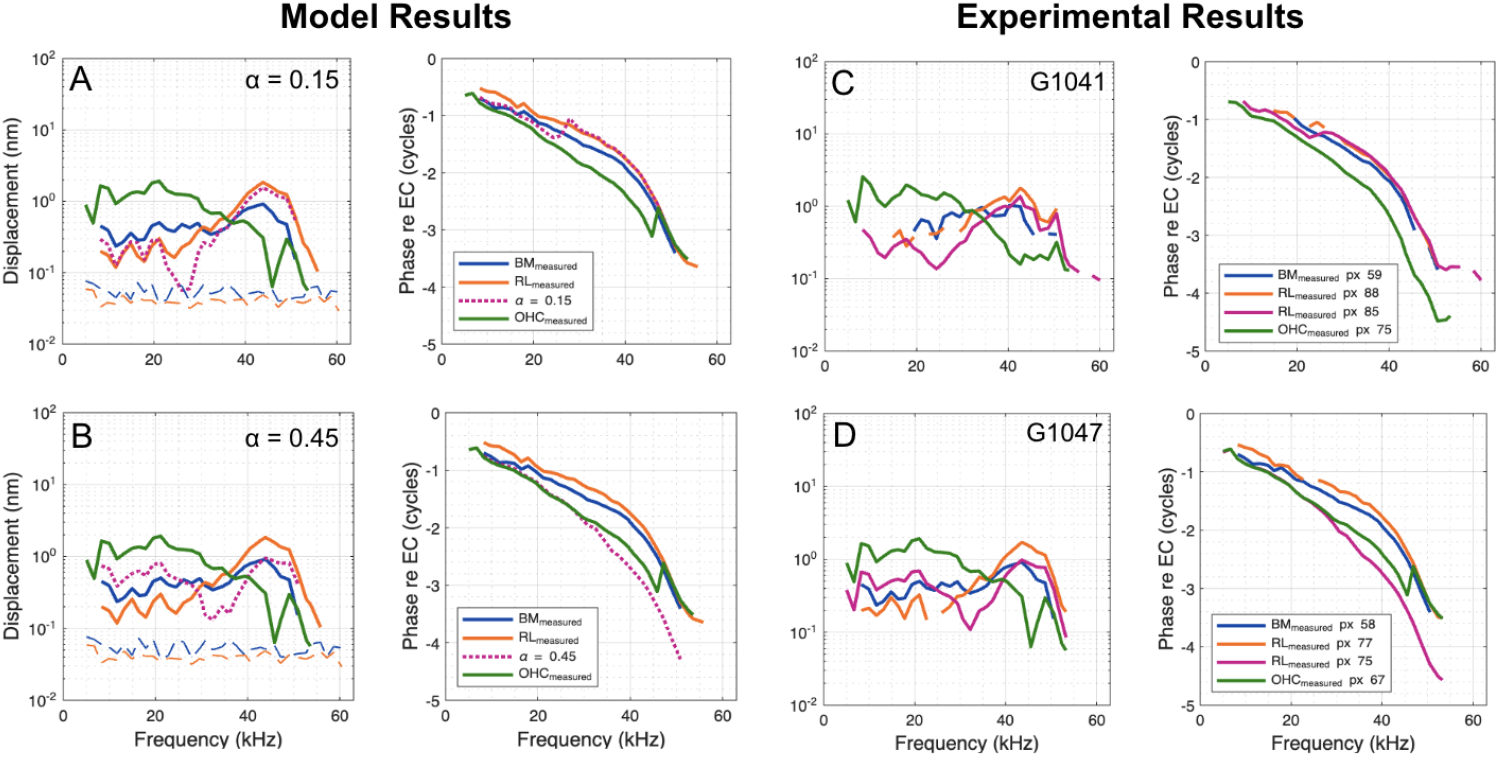
Comparison of 70 dB SPL experimental and model-predicted frequency responses with various α values. (A&B) Measured BM (pixel 58, blue), OHC/DC-region (pixel 67, green), and RL-region (pixel 77, orange) displacement and phase from G1047 (Fig. 3) along with model predictions of intra-RL motion (dotted pink) when (A) α = 0.15 and (B) α = 0.45. (C&D) Experimental results from G1041 (E) (also shown in Fig. 11A&B) and G1047 (F) (also shown in Fig. 4A&B) to compare to the model predicted results shown in (A&B).

### C. OHC/DC region

In measurements made through the RW at the 25 kHz BF location, Cooper et al. (2018) observed an abrupt increase in motion from the BM to the OHC/DC region (their Fig. 6). They surmised that the OHC/DC motion was elliptical, with a long axis running approximately perpendicular to the long axis of the DC phalangeal processes. Frost et al., (2025) decomposed the motion with measurements from two angles (at 70 and 80 dB SPL), and found the motion to be a high aspect-ratio ellipse (nearly lineal motion) with nearly equal longitudinal and transverse components (drawn as lineal motion in the green arrows in Fig. 16).

**Fig. 16:**
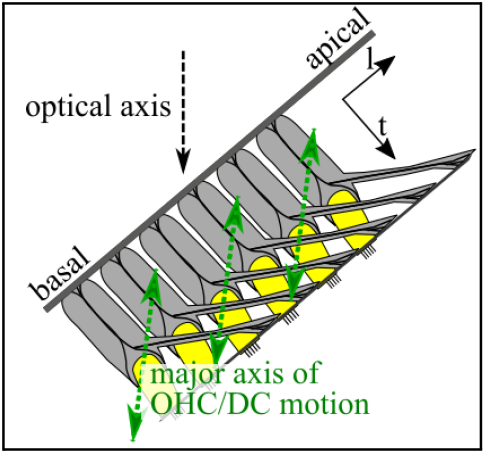
Cartoon diagram depiction of the orientation of the OCC in this work’s responses. The structures in the diagram are the DCs (grey), OHCs (yellow), BM, and RL. The optical axis of the OCT and major axis of OHC/DC motion is shown.

From the *l,r,t* values noted in the figure captions, when viewing through the RW to the 45 kHz place, our optical axis was primarily transverse but with a substantial negative longitudinal component (Fig. 16). With this angle of observation, a motion perpendicular to the longitudinal axis of the phalangeal processes will be approximately in line with the optical axis, and thus we are well-positioned to detect that motion. As noted above, the OHC/DC-region motion we observed extended to pixels that would be beyond (shallower than) the OHC base. The extensive and uniform OHC/DC motion we observed likely includes the nuclear region of OHCs and DCs, swinging perpendicular to the phalangeal processes.

Another noteworthy aspect of the OHC/DC motion is the phase, which, from low frequency to the BF, smoothly accumulates about 0.75 cycle phase lag relative to the BM (41). This is most clearly presented in Figs. 2C, 4B, 8B, and 11B, where OHC/DC and BM phases are plotted together. The steeper phase-versus-frequency slope indicates that the traveling wave motion is slower in the OHC/DC region than on the BM and the RL. Because the RL leads the BM motion by close to 0.25 cycle in the BF peak, the OHC/DC and RL regions are nearly in phase there.

### D. Conclusions

In conceptual models of cochlear operation, the longitudinal/transverse OHC/DC motion has been proposed to be central to cochlear amplification (40–42), but detailed cochlear models along those lines have not been developed. Another possibility is that the OHC/DC motion is a by-product of cochlear amplification, not central to its operation (43); it is distant from the RL (where MET occurs) and distant from the BM (supporting the cochlear traveling wave). This anatomically-promoted, diagonally swinging motion might represent uncoupling of the electromotile motion of the OHC base from the RL, allowing strains at the RL side of the OHC to provide the essential active force of the cochlear amplifier. Recent cochlear models (44) and concepts (45) employ active forcing at the RL.

In our observations, OHC electromotile strains were large in the transition between the RL and OHC/DC regions. In the BF peak, the region of strain generally extended through a wider distance at 70 than at 50 dB. Thus, the OHC/DC and RL region motions were less coupled at 50 than at 70 dB, and since lower SPL motions show relatively more BF-peak amplification, uncoupling appears to be advantageous to functional cochlear amplification. This line of reasoning seems to flow more with the second concept of cochlear amplification noted above – that active force is delivered at the RL, and the OHC/DC motion is not the essential driver of amplification – but physics-based modeling is needed to make any substantive claims. The experimental observations of the present work guide and constrain our understanding of the operation of the cochlear amplifier, providing fuel for conceptually-developed, then physics-based cochlear models.

## V. Data and Code Availability

Data and analysis code will be made available upon reasonable request.

## VI. Author Contributions

K.H.W. gathered and analyzed the data and co-wrote the manuscript. E.S.O. designed the research and co-wrote the manuscript. C.E.S. gathered the data, coded the initial analysis steps, and edited the manuscript.

## VII. Acknowledgements

We thank Lauren A. Chiriboga for commenting on the manuscript. This work was funded by the National Institute on Deafness and Other Communication Disorders grant DC015362.

## Notes

### Competing Interest Statement

The authors have declared no competing interest.

