## Supplementary Figures and Table. for "Active strains in the basal organ of Corti in gerbil"

### Supplemental Information

#### A. DPOAEs

DPOAEs were taken at the beginning of the experiment to assess cochlear health. The average DPOAEs are presented in Fig. S1.

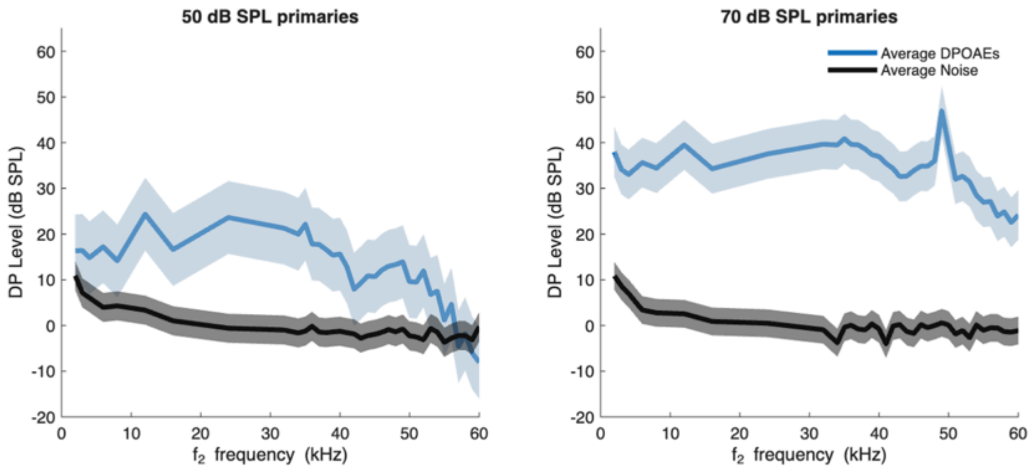


**Fig. S1.** Average initial DPOAEs (blue) of preparations 986, 1025, 1041, 1047, and 1049 and noise floor (grey). Dark line indicates the mean. The shaded area represents ± one standard deviation from the mean.

#### B. Displacement frequency responses for G986

The trends observed in the displacement frequency responses at 50 and 70 dB for G986 in Fig. S2 is consistent with what was observed for the other experiments.

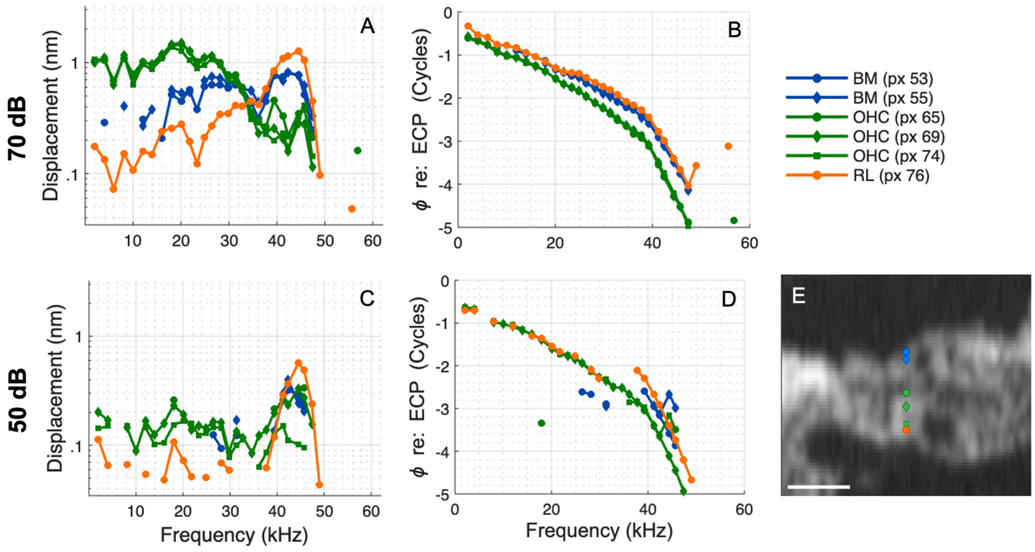


**Fig. S2.** Displacement frequency responses from 50 and 70 dB from seven locations along a single A-scan. (A&C) BM (blue), OHC-region (green), intra-RL-region (dashed orange), and RL-region (orange) displacements at 70 dB SPL (A) and 50 dB SPL (C). (B&D) The respective phase responses re: ECP. (E) B-scan with BM, OHC-region, intra-RL-region, and RL-region locations of measurements reported in (A-D) identified. Scale bar = 50 µm. Gerbil 986 Run 4. BF = 44.5 kHz.

**C. Strain frequency responses for G1025**

Fig. S3 displays the strain amplitude and phase calculated from pixels near the RL, 89-87, 87-86, and 86-85 are shown (C, D), along with the displacement data (A, B). There is a mild peak in the strain at ~30 kHz (C). The 89-87 and 86-85 strain phases are similar to each other throughout the frequency range (D). Below 30 kHz, the strain phase from the 87-86 calculation diverges from the others. The strain phase calculated from 89-87, when copied to panel B as a blue dashed line, aligns with the displacement phase of pixel 89 (closest to the RL surface) from 22 to 50 kHz. Below 22 kHz, the strain leads pixel 89 by ~0.2 cycle, and is ~1/2 cycle off from displacement pixels 85, 86, 87, similar to observations in Fig. 6B.

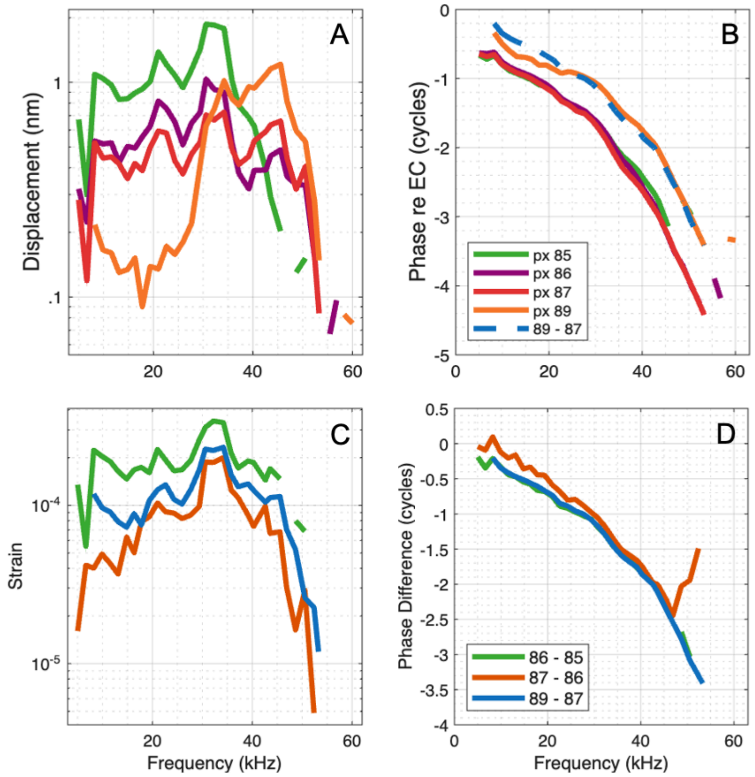


**Fig. S3.** Strain and displacement responses close to the RL at 70 dB. (A&B) The displacement amplitude and phase re: ECP of four pixels close to the RL. (C&D) The strain amplitude and strain phase calculated between the pixels in (A&B). The strain phase found with pixels 89 and 87 is plotted in (B) as a blue dashed line for comparison. Gerbil 1025 Run 13. BF = 45.5 kHz.

**D. G1049 Results**

The results from G1049 (Fig. S4-S6) were generally consistent with the results presented from the other experiments.

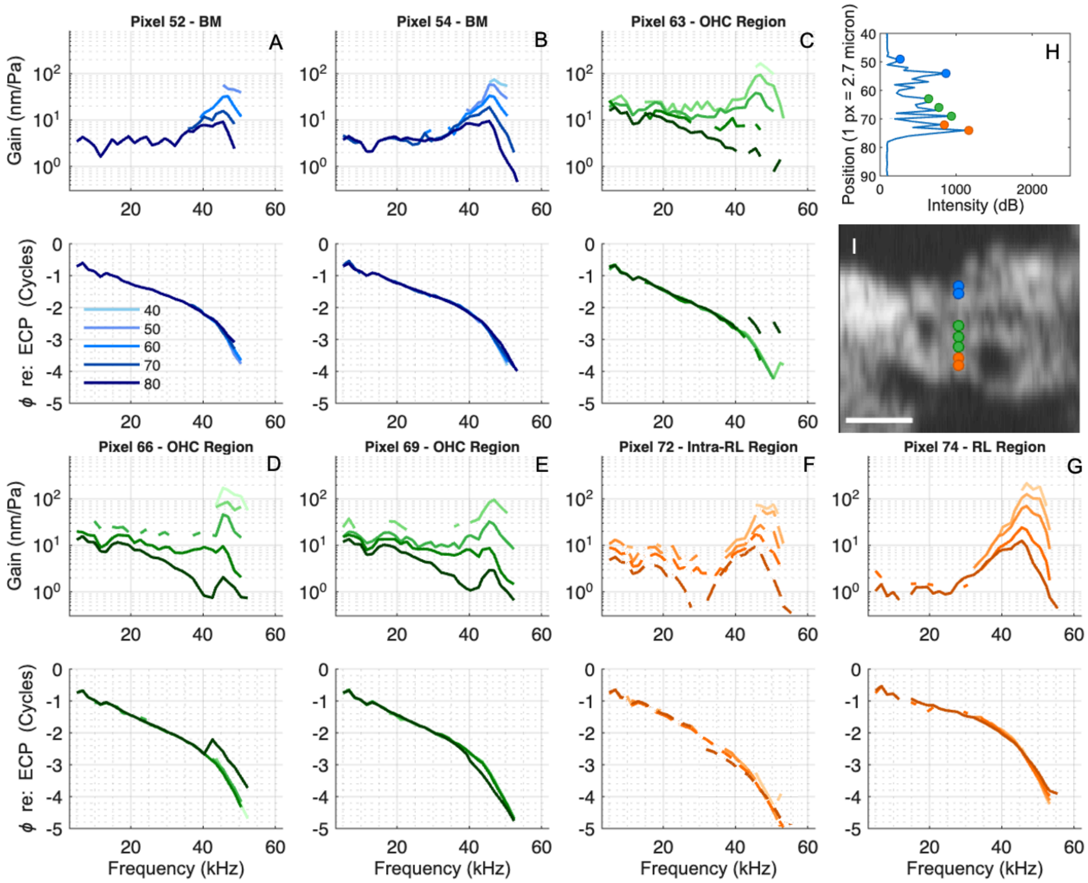


**Fig. S4.** Responses from seven locations along a single A-scan. (A-G) BM (blue), OHC-region (green), intra-RL-region (dashed orange), and RL-region (orange) gain. The respective phase responses of (A-G) re: ECP are plotted below the gain responses. (H) A-scan of 70 dB response. (I) B-scan with BM (blue), OHC-region (green), and intra-RL-region and RL-region (orange), locations of measurements reported in (A-G) are denoted with markers. Scale bar = 50 µm. Optical axis components were (*l*, *r*, *t*) = (-0.64, -0.03, 0.77). Gerbil 1049 Run 7. BF = 45.5 kHz.

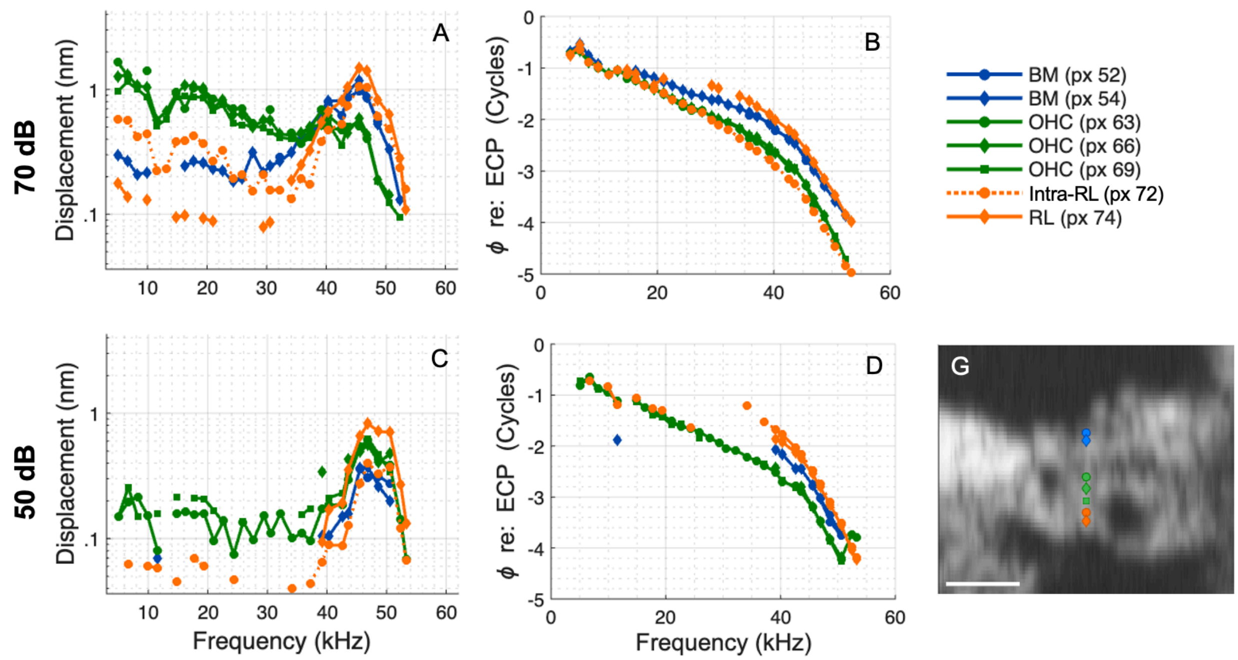


**Fig. S5.** Displacements and phases from seven locations along a single A-scan. (A&C) BM (blue), OHC-region (green), intra-RL-region (dashed orange), and RL-regions (orange) displacements at 70 dB SPL (A) and 50 dB SPL (C). (B&D) The respective phase responses re: ECP. (E) B-scan with BM (blue), OHC region (green), and intra-RL region and RL region (orange) locations of measurements reported in (A-D) identified. Scale bar = 50 µm. Gerbil 1049 Run 7. BF = 45.5 kHz.

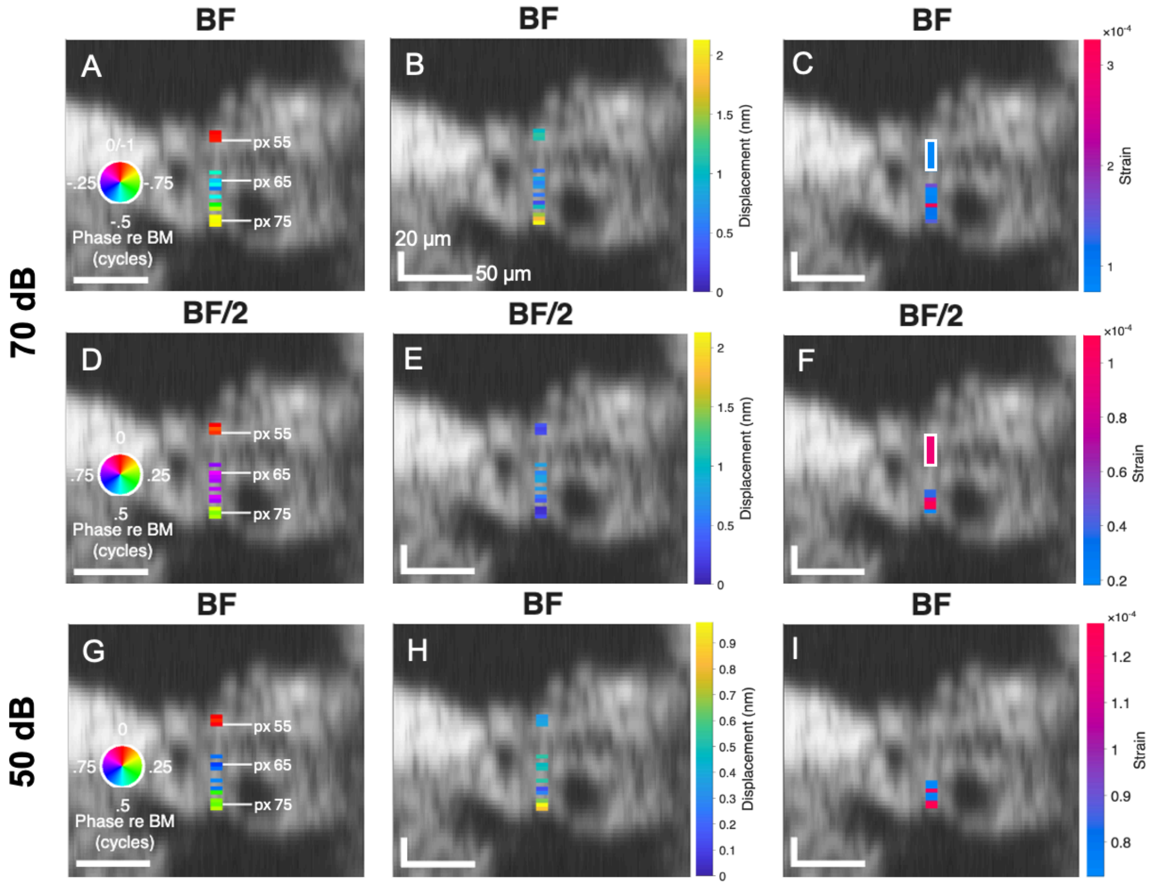


**Fig. S6.** Displacement amplitude and phase at all pixels deemed out of the noise along one A-scan, plotted as a 1D heat map onto the B-scan. (A, D, G) Displacement phase re BM: BF phase re: BM at 70 (A) and 50 dB (G); (D) BF/2 phase re: BM at 70 dB. (Pixel 53 was used for BM reference.) (B, E, H) Displacement at BF at 70 (B) and 50 dB (H), and at BF/2 at 70 dB (E). (C, F, I) Displacement strain at BF at 70 (C) and 50 dB (I), and at BF/2 at 70 dB (F). The white rectangle in (C&F) denotes a region where the strain was calculated over more than 3 pixels. Vertical scale bars = 20 µm and horizontal scale bars = 50 µm. Gerbil 1049 Run 7. BF = 45.5 kHz.

### E. G1041 alpha modeling

Fig. S7 demonstrates the modeling of the intra-region motion from the data taken from G1041. In Fig. S7B, with an α = 0.26, this replicates the phase crossing behavior of the RL region at a sub-BF frequency that is measured and observed in pixel 85 (refer to Fig. 11 and Fig. 12). By α = 0.35 (Fig. S7D), the x(α) behavior looks similar to the intra-RL region responses at pixel 83. There is a notch in the displacement response and a lag in the phase for a frequency region sub-BF, but then eventually becomes in-phase with the OHC/DC region. This notch appears more similar to pixel 83 at α = 0.45.

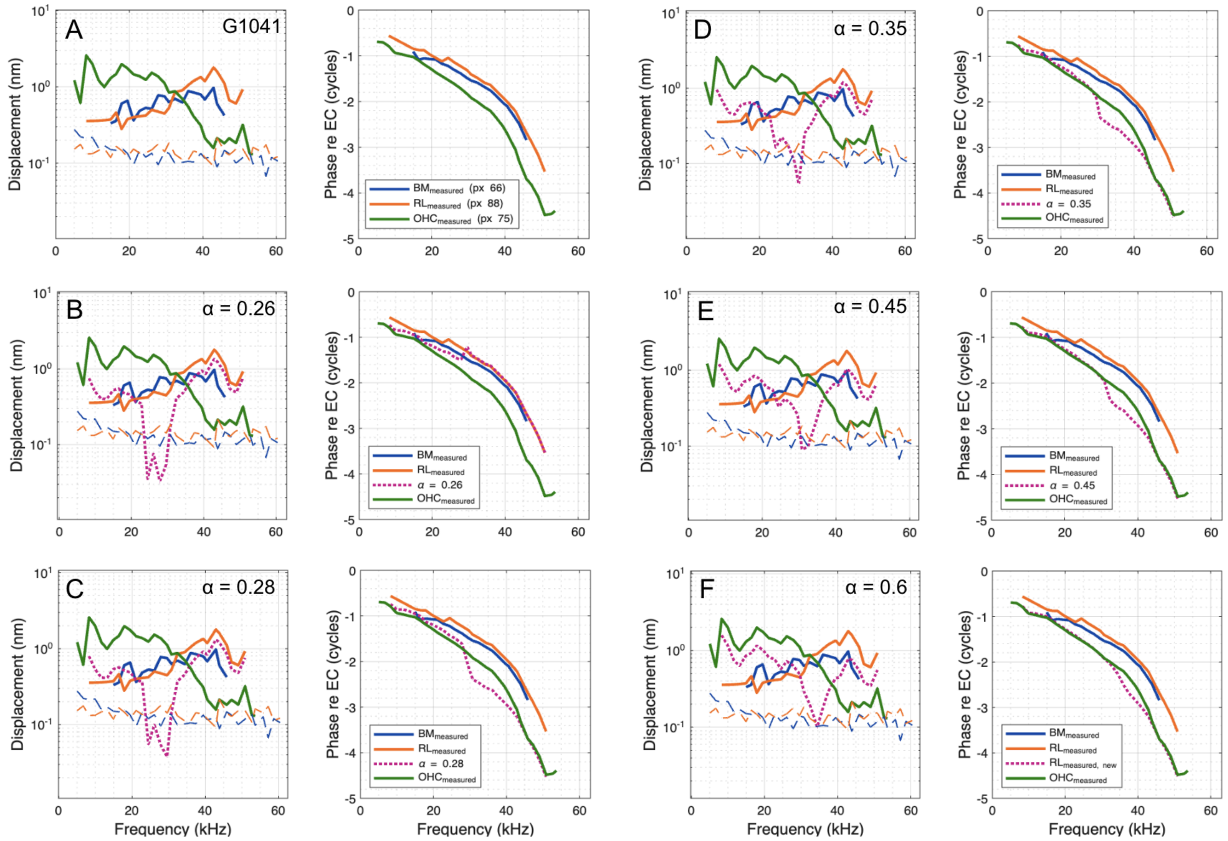


**Fig. S7.** (A) Measured BM (blue),OHC/DC-region (green), and RL-region (orange) motion of pixels 66, 75, and 88 at 70 dB SPL from G1041 (Fig. 11) used to model x(α) motion (dotted pink), when (B) α = 0.26, (C) α = 0.28, (D) α = 0.35, (E) α = 0.45, and (F) α = 0.6.

**F. Table 1**

Table 1 presents similarity percentages, strain values at the BF for 50 and 70 dB, and strain values at the BF/2 for 70 dB for G1047, G1025, G1041, G986, and G1049. These values were all calculated between a shallower and deeper pixel. If the similarity percentage was 70% or greater (refer to Methods) at 70 dB, the responses from the two pixels were deemed from the same region, and color-coded similarly in the figures. If the similarity percentage was below this cut-off, the responses were denoted as transitional between regions, "intra-region." The 70% cut-off was relaxed for a few color-coded curves, these are marked by an asterisk.

| **SPL:** | **70 dB** | | | | | **50 dB** | | | |
|---|---|---|---|---|---|---|---|---|---|
| **Gerbil** | **Shallower Pixel** | **Deeper Pixel** | **Similarity** | **Strain at BF** | **Strain at BF/2** | **Shallower Pixel** | **Deeper Pixel** | **Similarity** | **Strain at BF** |
| **1047** | 47 | 48 | 90% | | | 48 | 49 | 76% | |
| | 48 | 49 | 77% | | | 49 | 58 | 63% | |
| | 49 | 52 | 75% | | | 58 | 63 | -14% | 4.63E-05 |
| | 52 | 57 | 80% | | | 63 | 66 | 66% | |
| | 57 | 58 | 77% | 3.87E-05 | 4.74E-05 | 66 | 67 | 87% | |
| | 58 | 59 | 71% | 3.35E-05 | 1.03E-04 | 67 | 68 | 80% | |
| | 59 | 62 | -10% | 5.56E-05 | 2.27E-04 | 68 | 70 | 70% | |
| | 62 | 63 | 81% | | | 70 | 71 | 41% | |
| | 63 | 66 | 66%* | 4.72E-05 | 5.08E-05 | 71 | 73 | 54% | 2.52E-05 |
| | 66 | 67 | 95% | 1.23E-05 | 3.16E-05 | 73 | 75 | 8% | 1.19E-04 |
| | 67 | 68 | 94% | 4.53E-05 | 2.09E-05 | 75 | 76 | 80% | 6.02E-05 |
| | 68 | 70 | 90% | 1.51E-05 | 2.92E-05 | 76 | 77 | 85% | 3.33E-05 |
| | 70 | 71 | 93% | 1.65E-05 | 3.09E-05 | 77 | 78 | 83% | 3.99E-05 |
| | 71 | 73 | 69%* | 4.20E-05 | 7.06E-05 | | | | |
| | 73 | 75 | 9% | 1.19E-04 | 1.65E-04 | | | | |
| | 75 | 76 | 65% | 9.43E-05 | 9.07E-05 | | | | |

| | | | | | | | | | |
|---|---|---|---|---|---|---|---|---|---|
| | 76 | 77 | 34% | 1.46E-04 | 2.20E-04 | | | | |
| | 77 | 78 | 86% | 3.27E-05 | 2.54E-05 | | | | |
| **Gerbil** | **Shallower Pixel** | **Deeper Pixel** | **Similarity** | **Strain at BF** | **Strain at BF/2** | **Shallower Pixel** | **Deeper Pixel** | **Similarity** | **Strain at BF** |
| **1025** | 64 | 65 | 88% | | | 64 | 65 | 75% | |
| | 65 | 66 | 93% | | | 65 | 66 | 77% | |
| | 66 | 68 | 88% | | | 66 | 68 | 65% | |
| | 68 | 69 | 85% | | | 68 | 69 | 79% | |
| | 69 | 70 | 81% | | | 69 | 70 | 67% | |
| | 70 | 71 | 73% | 8.26E-05 | 2.79E-05 | 70 | 71 | 60% | 4.63E-05 |
| | 71 | 78 | -62% | 5.95E-05 | 1.04E-04 | 71 | 78 | -30% | |
| | 78 | 79 | 79% | 5.94E-05 | 7.80E-05 | 78 | 79 | 72% | |
| | 79 | 80 | 94% | 3.05E-05 | 3.36E-05 | 79 | 80 | 88% | |
| | 80 | 81 | 97% | 3.84E-05 | 6.48E-06 | 80 | 81 | 91% | 3.25E-05 |
| | 81 | 82 | 92% | 3.77E-05 | 4.11E-05 | 81 | 82 | 94% | |
| | 82 | 83 | 74% | 1.51E-04 | 1.17E-04 | 82 | 83 | 83% | 3.50E-05 |
| | 83 | 84 | 68% | 9.09E-05 | 1.52E-04 | 83 | 84 | 70% | |
| | 84 | 85 | 78% | 7.39E-05 | 1.17E-04 | 84 | 85 | 67% | 7.60E-05 |
| | 85 | 86 | 33% | 1.47E-04 | 1.93E-04 | 85 | 86 | -9% | 3.40E-04 |
| | 86 | 87 | 58% | 6.81E-05 | 9.10E-05 | 86 | 88 | 56% | 6.34E-05 |
| | 87 | 88 | 19% | 2.07E-04 | | 88 | 89 | 71% | 6.62E-05 |
| | 87 | 89 | -25% | | 1.35E-04 | | | | |
| | 88 | 89 | 82% | | | | | | |

| Gerbil | Shallower Pixel | Deeper Pixel | Similarity | Strain at BF | Strain at BF/2 | Shallower Pixel | Deeper Pixel | Similarity | Strain at BF |
|---|---|---|---|---|---|---|---|---|---|
| 1041 | 56 | 59 | 73% | | | 66 | 75 | 30% | |
| | 59 | 60 | 81% | | | 75 | 76 | 71% | 8.46E-05 |
| | 60 | 63 | 78% | | | 76 | 77 | 79% | 5.59E-05 |
| | 63 | 64 | 77% | | | 77 | 78 | 70% | 6.61E-05 |
| | 64 | 66 | 66%* | | | 78 | 80 | 52% | 3.98E-05 |
| | 66 | 70 | 37% | 1.76E-05 | 6.37E-05 | 80 | 81 | 34% | |
| | 70 | 75 | 60% | 4.02E-06 | 1.58E-05 | 81 | 82 | 38% | |
| | 75 | 76 | 87% | | 4.44E-05 | 82 | 83 | 73% | 3.80E-05 |
| | 76 | 77 | 92% | | | 83 | 84 | 66% | 4.91E-05 |
| | 76 | 80 | 9% | 3.23E-05 | | 84 | 85 | 47% | 1.07E-04 |
| | 77 | 78 | 70% | | 1.77E-04 | 85 | 88 | 79% | |
| | 78 | 80 | 34% | | 7.79E-05 | | | | |
| | 80 | 81 | 72% | 1.20E-04 | | | | | |
| | 81 | 82 | 67% | 9.96E-05 | 3.43E-05 | | | | |
| | 82 | 83 | 73% | 9.46E-05 | 6.00E-05 | | | | |
| | 83 | 84 | 56% | 7.83E-05 | 8.81E-05 | | | | |
| | 84 | 85 | 81% | 6.98E-05 | 2.06E-05 | | | | |
| | 85 | 88 | 57%* | 7.54E-05 | 4.73E-05 | | | | |
| **Gerbil** | **Shallower Pixel** | **Deeper Pixel** | **Similarity** | **Strain at BF** | **Strain at BF/2** | **Shallower Pixel** | **Deeper Pixel** | **Similarity** | **Strain at BF** |
| **986** | 51 | 53 | 78% | 2.19E-05 | 2.91E-05 | 51 | 53 | 81% | |

| | | | | | | | | | |
|---|---|---|---|---|---|---|---|---|---|
| | 53 | 55 | 80% | | | 53 | 55 | 66% | |
| | 55 | 60 | 72% | 1.95E-05 | 1.24E-05 | 55 | 60 | 46% | |
| | 60 | 65 | -54% | 5.22E-05 | 1.03E-04 | 60 | 69 | 27% | 2.43E-05 |
| | 65 | 69 | 89% | 6.02E-06 | | 69 | 74 | 57% | 1.73E-05 |
| | 69 | 74 | 86% | 1.10E-05 | 1.66E-05 | 74 | 76 | -3% | 1.08E-04 |
| | 74 | 76 | -75% | 2.21E-04 | 2.06E-04 | | | | |
| **Gerbil** | **Shallower Pixel** | **Deeper Pixel** | **Similarity** | **Strain at BF** | **Strain at BF/2** | **Shallower Pixel** | **Deeper Pixel** | **Similarity** | **Strain at BF** |
| **1049** | 53 | 54 | 75% | | | 53 | 54 | 86% | |
| | 54 | 55 | 85% | | | 54 | 55 | 90% | |
| | 55 | 63 | -38% | 7.44E-05 | 9.55E-05 | 55 | 63 | 21% | |
| | 63 | 65 | 62%* | | | 63 | 65 | 74% | |
| | 65 | 66 | 82% | | | 65 | 66 | 79% | |
| | 66 | 67 | 68%* | 1.61E-04 | | 66 | 69 | 80% | |
| | 67 | 69 | 70% | 7.90E-05 | | 69 | 71 | 34% | 7.26E-05 |
| | 69 | 71 | 62% | 8.37E-05 | 3.77E-05 | 71 | 72 | 1% | 1.24E-04 |
| | 71 | 72 | 18% | 3.25E-04 | 9.14E-05 | 72 | 74 | 24% | 7.48E-05 |
| | 72 | 74 | 39% | 1.13E-04 | 1.10E-04 | 74 | 75 | 62% | 1.25E-04 |
| | 74 | 75 | 80% | 1.07E-04 | 1.83E-05 | 75 | 76 | 69% | 1.27E-04 |
| | 75 | 76 | 69% | 1.41E-04 | | | | | |
